# Distinct neuronal activity patterns induce different gene expression programs

**DOI:** 10.1101/146282

**Authors:** Kelsey M. Tyssowski, Ramendra N. Saha, Nicholas R. DeStefino, Jin-Hyung Cho, Richard D. Jones, Sarah M. Chang, Palmyra Romeo, Mary K. Wurzelmann, James M. Ward, Serena M. Dudek, Jesse M. Gray

**Author notes:** These authors contributed equally (Kelsey M. Tyssowski, Ramendra N. Saha, Nicholas R. DeStefino). These authors contributed equally, co-corresponding authors (J.M.G.), (S.M.D). Lead contact: Jesse Gray, 77 Ave Louis Pasteur, Boston, MA, 02115.

## Abstract

Brief and sustained neuronal activity patterns can have opposite effects on synaptic strength that both require activity-regulated gene (ARG) expression. However, whether distinct patterns of activity induce different sets of ARGs is unknown. In genome-scale experiments, we reveal that a neuron’s activity-pattern history can be predicted from the ARGs it expresses. Surprisingly, brief activity selectively induces a small subset of the ARG program that that corresponds precisely to the first of three temporal waves of genes induced by sustained activity. These first-wave genes are distinguished by an open chromatin state, proximity to rapidly activated enhancers, and a requirement for MAPK/ERK signaling for their induction. MAPK/ERK mediates rapid RNA polymerase recruitment to promoters, as well as enhancer RNA induction but not histone acetylation at enhancers. Thus, the same mechanisms that establish the multi-wave temporal structure of ARG induction also enable different sets of genes to be induced by distinct activity patterns.

## INTRODUCTION

Neurons induce expression of hundreds of activity-regulated genes (ARGs) in response to elevations of neuronal activity (Flavell and Greenberg, 2008). This gene induction is required for many forms of long-lasting synaptic plasticity, including, intriguingly, both synaptic strengthening and weakening. Brief, high frequency activity strengthens synapses through LTP (Abbott and Nelson, 2000), whereas sustained activity weakens synapses through homeostatic scaling (Turrigiano, 2008). The fact that LTP and homeostatic scaling both require activity-regulated transcription (Ibata et al., 2008; Nguyen et al., 1994) suggests that different patterns of neuronal activity may induce sets of ARGs with opposite functions. Consistent with this idea, different patterns of neuronal activity differentially induce the expression of several individual ARGs (Douglas et al., 1988; Greenberg et al., 1986; Sheng et al., 1993; Worley et al., 1993). Because induced mRNAs remain in the cell for hours to days (Schwanhausser et al., 2011), these observations hint at the possibility that a neuron’s activity pattern history might be predictable from its gene expression state (*i.e.*, set of expressed genes). Given that there are hundreds of ARGs, neurons have the potential to encode a vast number of activity patterns in gene expression states. However, the lack of genome-scale comparisons of ARG expression between even two distinct activity patterns has precluded an understanding of how neuronal activity patterns are coupled to ARG expression.

The ARG expression program is likely structured temporally into two major waves of gene induction that are well characterized in non-neuronal cells (Fowler et al., 2011; Herschman, 1991). The first of these waves comprises primary response genes (PRGs, often called immediate-early genes), which do not require de novo translation for their induction. The second major wave of gene induction comprises secondary response genes (SRGs), which require de novo translation for their induction and are regulated by PRG protein products (Fowler et al., 2011; Herschman, 1991; Yamamoto and Alberts, 1976). Given that neurons also induce at least two temporally distinct waves of transcription (Flavell and Greenberg, 2008; West and Greenberg, 2011), the ARG program is likely also divided into PRGs and SRGs. However, it is not known how this multi-wave structure relates to the ability of the ARG program to distinguish different activity patterns.

Neuronal activity induces ARGs via calcium influx and activation of calcium-dependent signaling pathways, but it is not known how these signaling pathways mediate activity-pattern-specific gene expression. The many calcium-dependent signaling pathways and hundreds of associated signaling molecules (Fields et al., 2005; Flavell and Greenberg, 2008) provide, in principle, enough complexity to couple many distinct neuronal activity patterns to corresponding gene expression states. Indeed, calcium-dependent signaling pathways respond differentially to different patterns of calcium influx as well as to different patterns of neuronal activity (De Koninck and Schulman, 1998; Dolmetsch et al., 1998, 1997; Dudek and Fields, 2001; Eshete and Fields, 2001; Fields et al., 1997; Fujii et al., 2013; Ma et al., 2011; Wu et al., 2001). An intriguing possibility for how different calcium-dependent signaling dynamics might induce distinct sets of genes is suggested by experiments in cancer cell lines. Brief signaling pathway activation is sufficient to induce PRGs, but sustained signaling pathway activation is required to stabilize PRG protein products, presumably allowing them to persist for long enough to induce SRGs (Murphy et al., 2004, 2002). In this way, signaling pathway activation and the temporal structure of ARG transcription could intersect to couple distinct activity patterns to expression of different subsets of genes.

We performed a genome-scale comparison of ARG induction in response to two neuronal activity patterns: brief activity and sustained activity. Using this simplified model, we were able to predict the duration of a neuron’s elevated activity from its ARG gene expression state, a formal demonstration that specific patterns of neuronal activity are indeed encoded in ARG expression. As expected, sustained activity induces both PRGs and SRGs. Surprisingly, we found that brief activity induces only a small subset of PRGs, rapid PRGs, which we show correspond to the first of three distinct temporal waves of ARG induction. Rapid PRGs are distinguished from genes in subsequent waves by a requirement for MAPK/ERK signaling for their induction. Abolishing MAPK/ERK signaling blunts and delays rapid PRG induction, altering the temporal structure of the ARG program. It also blocks gene induction in response to brief activity. Thus, a signaling mechanism that helps create the multi-wave temporal structure of the ARG program also enables distinct activity patterns to generate different gene expression states.

## RESULTS

### Rapid but not delayed PRGs are induced by brief activity

We investigated the possibility that neuronal activity pattern histories are encoded in ARG expression states by focusing on two activity patterns: brief and sustained activity. To determine whether brief and sustained neuronal activation induce different subsets of genes, we activated neurons either briefly (1-5 minutes) or continuously (up to six hours) (Figure 1A). Our primary method of neuronal activation was KCl-mediated membrane depolarization of mouse cortical neurons. This method enables cell-to-cell consistency and control over activity patterns that are difficult to achieve in vivo. Sustained KCl-mediated depolarization elevates intraceullular calcium for a minimum of 20 minutes, but likely indefinitely (Dolmetsch et al., 2001; Evans et al., 2013). In contrast, brief KCl-mediated depolarization elevates intracellular calcium only during the period of elevated KCl (Kingsbury et al., 2007). To ensure that our observations would not be unique to a specific stimulation method or mammalian species, we also used bicuculline-mediated synaptic stimulation of rat cortical neurons. Sustained bicuculline/4AP treatment maintains an elevated firing rate of neurons in culture for at least twenty minutes (Arnold et al., 2005; Li et al., 2007). To elicit a brief increase in neuronal firing rate, we added a voltage-gated sodium channel blocker (TTX) shortly after bicuculline/4AP stimulation.

**Figure 1.**
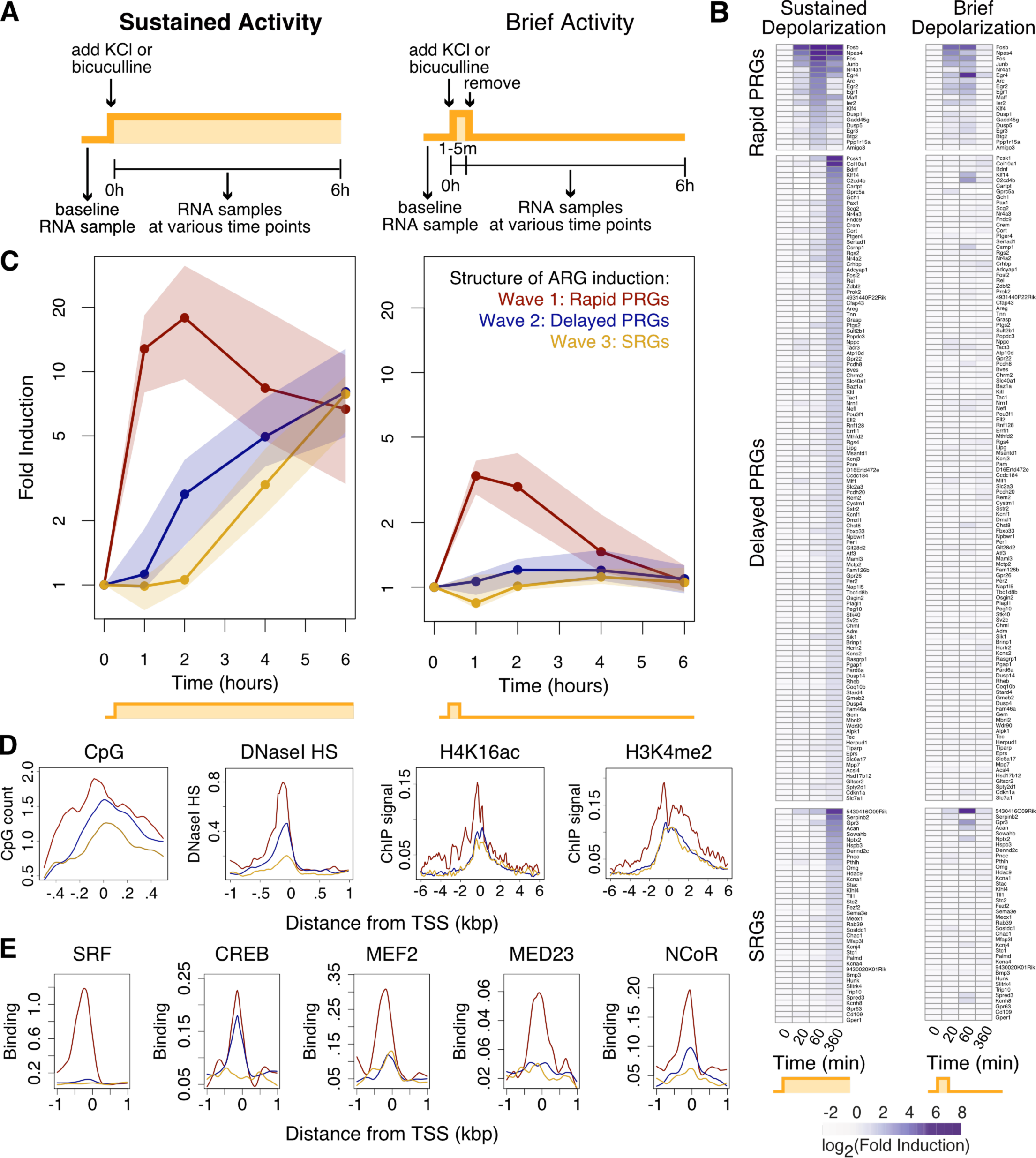
Brief neuronal activation selectively induces the first of three waves of gene induction. (A) Experimental system for comparing sustained and brief neuronal activation in vitro. The yellow stimulation diagram is used throughout the figures to indicate sustained or brief activity. Except where indicated otherwise, neuronal activation is accomplished with brief (1 min) or sustained KCl-depolarization of DIV7-8 cortical neurons (from E16.5 mice) silenced 14-16h before stimulation with APV and NBQX. (B) Comparison of gene induction upon sustained or brief neuronal activation using activity-regulated-gene-capture-based RNA sequencing (ARG-seq). Shown are means from n=3-6 biological replicates. Genes shown are all induced >2 fold with FDR<0.05 (at any time point, with either brief or sustained stimulation). Gene categories are defined based on kinetics of neuronal activity-induced gene induction, as well as induction in the presence or absence of the translation-inhibitor cycloheximide (Figure S1B). PRG = primary response gene. SRG = secondary response gene. Genes induced by brief neuronal activation are enriched for rapid genes (p < 10^˗13^, Fisher’s exact test). (C) Three kinetically distinct temporal waves of gene induction, consisting of rapid PRGs, delayed PRGs, and SRGs, as detected by high-throughput microfluidic qPCR on the Fluidigm platform. For each gene, average expression values were determined from n=6 biological replicates and plotted with dots and solid lines representing the median from each gene class and shading covering the middle quartiles (25%-75%). Each wave is kinetically distinct from the other waves (rapid PRG induction is higher than delayed PRG or SRG induction at 1h, delayed PRG induction is greater than SRG induction at 2h, p<0.003, rank-sum test). There are 15, 37, and 9 genes from waves 1-3 respectively. (D) Chromatin state in unstimulated neurons at rapid PRGs, delayed PRGs, and SRGs shown in metaplots of the geometric mean signal for all genes in each category. CpG content, DNaseI hypersensitivity, H4K16ac and H3K4me2 are significantly different between rapid PRGs and delayed PRGs or SRGs (p<0.009, rank sum test on the area under the curves shown). ChIP signal is input-normalized read density. H4K16ac and H3K4me2 ChIP-seq data are from Telese et al., 2015. DNasel hypersensitivity data is from the ENCODE Project (ENCODE Project Consortium et al., 2012). (E) Transcription factor binding in unstimulated neurons from ChIP-seq, shown in metaplots as in (D). SRF, MEF2, MED23, and NCoR are significantly different between rapid PRGs and delayed PRGs or SRGs (p<0.009, rank sum test on the area under the curves shown). CREB is not significantly different between rapid and delayed PRGs (p=0.2), but is different between rapid PRGs and SRGs (p=0.007, rank sum test). Binding is calculated from input-normalized ChIP-seq read density. Data from Kim et al., 2010; Telese et al., 2015. Related to figures S1 and S2.

We assessed ARG expression in a time course following activation using digital hybridization-based counting (NanoString) and two RNA-sequencing (RNA-seq) approaches. The RNA-seq approaches included total RNA-seq and targeted sequencing (Mercer et al., 2011) of 251 ARGs that previously showed >3.5 fold induction in response to KCl treatment (Kim et al., 2010) (Table S1). We confirmed that this targeted ARG-sequencing (ARG-seq) produced results that correlate with our total RNA-seq results (r^2^ = 0.9, Figure S1A). ARG-seq resulted in a ~76-fold enrichment of reads in targeted regions compared to total RNA-seq (Figure S1A). This allowed us to quantify ARG induction with as few as 3 million reads per sample, enabling us to perform RNA-seq from a total of 123 samples across all of the experiments in our study.

We used these methods to ask whether the ARGs induced by brief and sustained activity patterns differ in composition or dynamics. We hypothesized that PRGs but not SRGs would be induced by brief neuronal activity because of the time required to synthesize and stabilize the PRG protein products that regulate SRGs. Because PRGs and SRGs had not been previously defined based on de-novo-translation-dependence in neurons, we first classified ARGs as PRGs or SRGs by performing ARG-seq on neurons treated with sustained KCl in the presence of the translation inhibitor cycloheximide (CHX) (processed ARG-Seq data provided in Table S2). We observed significant induction of 173 high-confidence ARGs in response to membrane depolarization using ARG-seq (Figure 1B). The majority of these ARGs (135) show little CHX sensitivity (<2 fold change), and we therefore classified them as PRGs (Figure S1B). The remaining 38 show a >2 fold reduction in expression in the presence of CHX, and we thus classified them as SRGs. In agreement with our hypothesis, brief activity was not sufficient to induce SRGs (Figure 1B). Surprisingly, however, brief activity significantly induced only a small subset (15/135) of PRGs (FDR<0.05, fold change>1.4, Figure 1B), indicating that de-novo-translation-independence is not the only requirement for induction in response to brief activity. Moreover, the distribution of mRNA expression levels of the brief-activity-insensitive genes was not shifted in response to brief activity at any time point, suggesting that this failure to observe induction was not merely due to insufficient sensitivity to detect small changes in individual genes (Figure S1C). Given that brief and sustained activity induce different gene sets, we asked whether it would be possible to predict whether neurons had undergone brief or sustained activity solely by assessing their gene expression states. Indeed, a nearest-neighbor classifier was able to correctly identify samples as having been stimulated with brief or sustained KCl-mediated depolarization using normalized expression values from all significantly induced genes (100% correct identification) or all genes captured by ARG-seq (92% correct identification), but not constitutively active captured genes (50% correct identification). This high-confidence prediction indicates that a neuron’s activity history is indeed encoded in its ARG expression state.

We next asked what distinguishes the few PRGs induced by brief neuronal activity from the remaining PRGs. We hypothesized that the PRGs induced by brief activity might be those most rapidly induced by sustained activity. To address this hypothesis, we divided PRGs into rapid and delayed PRGs based on their relative induction at one and six hour time points, similar to a previous classification of inducible genes in human cell lines (Tullai et al., 2007). We found that only 13% of PRGs are rapid PRGs, defined as those that show greater induction following one compared to six hours of sustained activity. In contrast, 87% of PRGs are delayed PRGs, defined as those that have induction kinetics similar to SRGs, showing greater induction at six compared to one hour of sustained activity (Figure 1B). An unsupervised method, principal component analysis, showed clear separation between rapid PRGs and other ARGs (i.e., delayed PRGs and SRGs), reinforcing our classification (Figure S1D). Moreover, a higher resolution time course using high-throughput qPCR on the Fluidigm platform both validated our ARG-seq findings and revealed that delayed PRGs are induced after rapid PRGs but before SRGs (Figure 1C, S1E). Therefore, activity-dependent transcription in neurons in response to sustained activity consists of at least three kinetically and mechanistically distinct waves of gene induction: a first, de-novo-protein-synthesis-independent wave (rapid PRGs); a second, de-novo-protein-synthesis-independent wave (delayed PRGs), and a third, de-novo-protein-synthesis-dependent wave (SRGs). In addressing our hypothesis, we found impressively that rapid PRGs were much more likely to be induced by brief activity than delayed PRGs or SRGs (p < 10^˗13^, Fisher’s exact test), and 14 of the 15 genes significantly induced by brief activity are rapid PRGs. Thus, of the 19 rapid PRGs, only 5 are not significantly induced by brief activity, and four of these five exhibit a trend toward induction by brief activity (mean fold change >1.5, Figure 1B). Analysis of pre-mRNA expression using total RNA-seq reads aligning to introns, a proxy for transcriptional activity (Gaidatzis et al., 2015; Gray et al., 2014), recapitulated our findings with mRNA analysis (Figure S1G), suggesting that the differences in induction kinetics and responsiveness to brief activity that we observed between rapid and delayed PRGs are due to transcriptional rather than post-transcriptional mechanisms. We also confirmed that the selective induction of rapid but not delayed PRGs by brief activity is not specific to KCl-mediated depolarization or to mice, as it also occurs following brief bicuculline-induced activity in rat primary cortical neurons (Figure S1F). These findings suggest that rapid PRGs are distinguished from delayed PRGs by as-yet-unknown transcriptional mechanisms that allow them to respond both rapidly and to brief activity.

We next investigated whether the genes in each of the three waves of ARG induction differ in their known or annotated gene function. Most (14 of 19) rapid PRGs that we identified in mouse cortical neurons are involved in regulation of transcription (e.g. *Fos, Npas4, Egr1-4, Btg2*). Rapid PRGs are also more likely than delayed PRGs or SRGs to be stimulus-induced in macrophages (p = 0.0004, Fisher’s exact test) (Escoubet-Lozach et al., 2011) and human cancer cell lines (p=0.0001, Fisher’s exact test) (Tullai et al., 2007), consistent with the idea that transcription factors are re-used in many cell types. Of the five rapid PRGs that do not directly regulate transcription, three, *Ppp1r15a* (a.k.a. *Gadd34*), *Dusp5*, and *Dusp1*, are components of signaling pathways that could regulate either transcription or other activity-dependent cellular processes (Bollen et al., 2010; Kholodenko et al., 2010). Thus, only two of the rapid PRGs, *Amigo3* and *Arc*, have no known direct or indirect role in transcriptional regulation. *Amigo3*, mediates cell-to-cell interactions (Kuja-Panula et al., 2003), but is relatively weakly induced by activity. *Arc* is a well-characterized gene that mediates activity-dependent AMPA receptor recycling and synaptic plasticity (Chowdhury et al., 2006; Plath et al., 2006). In contrast to rapid PRGs, delayed PRGs include many effector (i.e. non-transcription factor) genes, including neurotrophins (e.g. *Bdnf, Nrn1*), G-protein coupled receptors (e.g. *Gprc5a, Chrm2*), potassium channels (e.g. *Kcnf1, Kcnj3, Kcns2*), cell adhesion molecules (e.g. *Bves, Pcdh20*), and transporters (e.g. *Slc6a17, Slc40a1*). SRGs are more similar in function to delayed PRGs than to rapid PRGs, and they also include several potassium channels (e.g. *Kcna1, Kcna4, Kcnh8, Kcnj4*) and G-protein coupled receptors (e.g. *Gper1, Gpr3, Gpr63*). Because most of the effector ARGs are not induced by brief activity, we predict that the transcription-dependent synaptic changes caused by brief activity are driven by the protein products of only a few genes, including *Arc*. Importantly, we found that brief activity induced ARC protein in a de-novo-transcription-dependent manner (Figure S1H), consistent with the idea that ARC could mediate the synaptic changes driven by brief activity. The induction of transcription factors could also play a functional role in responding to brief neuronal activity by priming the neuronal genome to respond more quickly to future activity.

### Rapid PRG promoters are distinguished by open, active chromatin and the presence of pre-bound transcription regulators

Given that rapid PRGs are unique in their abilities both to be induced rapidly and to be induced by brief activity, we investigated what might enable these abilities. We noticed that rapid PRGs have a median gene length that is ~13 kb shorter than delayed PRGs and SRGs (Figure S2A). While this difference could contribute to the rapidity of rapid PRG induction, it is unlikely to account for the delay in induction of delayed PRGs and SRGs, given that we observe no increase in transcription even at the first exons of delayed PRGs or SRGs at early time points (Figure S2B). We next hypothesized that an open chromatin state at rapid PRG promoters in unstimulated neurons might facilitate their rapid induction and ability to be induced in response to brief elevations in neuronal activity. We found that rapid PRG promoters, compared to the promoters of other ARGs, have higher CpG and GC content (Figure 1D, S2C), which is associated with nucleosome depletion and increased transcription (Fenouil et al., 2012). To investigate chromatin state more directly, we assessed DNaseI hypersensitivity, a measure of chromatin accessibility, using data from embryonic mouse whole brain and 8 week mouse cerebrum (ENCODE Project Consortium et al., 2012). We found that rapid PRGs and constitutively active genes have higher DNaseI hypersensitivity, indicative of more open chromatin, compared to delayed PRGs and SRGs (Figure 1D, S2E). We next assessed the presence of active chromatin marks at ARG promoters in unstimulated neurons, using chromatin immunoprecipitation sequencing (ChIP-seq) data from cultured cortical neurons (Kim et al., 2010; Telese et al., 2015). Compared to delayed PRGs and SRGs, we found that rapid PRG promoters have more of the active chromatin marks H4K16ac, H3K4me2, and H3K27ac (Figure 1D, S2C). These histone marks also extend across a wider promoter-proximal region and are more bimodal at rapid PRGs, indicative of reduced nucleosome occupancy at or near transcription start sites (TSSs). These results indicate that in unstimulated neurons, rapid PRG promoters are in a relatively open chromatin state and may therefore be poised for rapid activation in response to brief activity.

The relatively open chromatin state at rapid PRG promoters in unstimulated neurons prompted us to ask whether these promoters might be selectively pre-bound to transcriptional activators, co-activators, or transcription machinery prior to neuronal activation. Consistent with the idea that pre-bound, paused Pol2 can maintain an open chromatin state to facilitate rapid transcriptional activation (Gilchrist et al., 2010; Saha et al., 2011), we found that Pol2 occupancy in unstimulated neurons is higher at the promoters of rapid PRGs and constitutively active genes compared to delayed PRGs and SRGs, using both published (Kim et al., 2010) (Figure S2E) and newly generated (Figure S5E) data. We next used published ChIP-seq data from unstimulated cultured cortical neurons to examine binding of transcriptional activators and coactivators (Kim et al., 2010; Telese et al., 2015). We found that compared to delayed PRG and SRG promoters, rapid PRG promoters have greater binding of the neuronal activity-regulated transcription factors SRF and MEF2, as well as the Mediator subunits MED23 and MED1 (Figure 1E, S2D). In contrast, the transcription factor CREB is pre-bound to a similar extent to rapid and delayed PRGs, but is not pre-bound to SRGs (Figure 1E, S2D). However, delayed PRG and SRG promoters eventually become accessible to transcription factors, as the activity-inducible transcription factor NPAS4 is similarly recruited to promoters of all three gene classes upon two hours of neuronal activation (Figure S2D). Interestingly, despite the open chromatin and binding of transcriptional activators at rapid PRG promoters, these genes have less transcription in unstimulated neurons than constitutive genes and no more than delayed PRGs or SRGs (Figure S2E). It is therefore possible that rapid PRGs are actively repressed in unstimulated neurons. Indeed, we observed greater binding of the NCoR repressor complex at rapid PRG promoters compared to delayed PRG or SRG promoters (Figure 1E, S2E). These data suggest that in addition to an open chromatin state, pre-binding of Pol2, SRF, MEF2, and Mediator may uniquely poise rapid PRGs for rapid induction, possibly creating a need for NCoR-mediated repression in unstimulated conditions.

### The MAPK/ERK pathway is required for the first wave of gene induction

We next asked whether rapid PRGs could respond quickly and to brief activity because they are targeted by a signaling pathway with these same response properties. In evaluating this possibility, we compared rapid and delayed PRGs, but we excluded SRGs to eliminate the confounding possibility of altered PRG induction affecting SRG induction. We hypothesized that the MAPK/ERK pathway might mediate rapid PRG induction based on its functions in transcription-dependent late-phase LTP and learning (Thomas and Huganir, 2004), which can be driven by brief neuronal activation. We first assessed MAPK/ERK pathway activation by western blotting for the pathway’s terminal kinase, phospho-ERK (pERK). We found that in response to both brief and sustained activity, pERK levels reached the same peak magnitude by five minutes after the start of the stimulus (Figures 2A-D), suggesting that the MAPK/ERK pathway is fully activated by brief activity, at least in the cytoplasm. Because many of the mechanisms by which MAPK/ERK signaling activates transcription require it to be active in the nucleus (Thomas and Huganir, 2004), we assessed nuclear pERK. We observed pERK the nucleus within 2 minutes of the initiation of either brief or sustained neuronal activity (Figure 2E), suggestive of rapid nuclear MAPK/ERK activation. Interestingly, upon brief stimulation, ERK activity in both the nucleus and the cytoplasm remains elevated for at least fifteen minutes after the removal of stimulus. This persistence of ERK activation suggests that MAPK/ERK pathway activity may retain the memory of brief activity long enough to activate the transcription of rapid PRGs.

**Figure 2.**
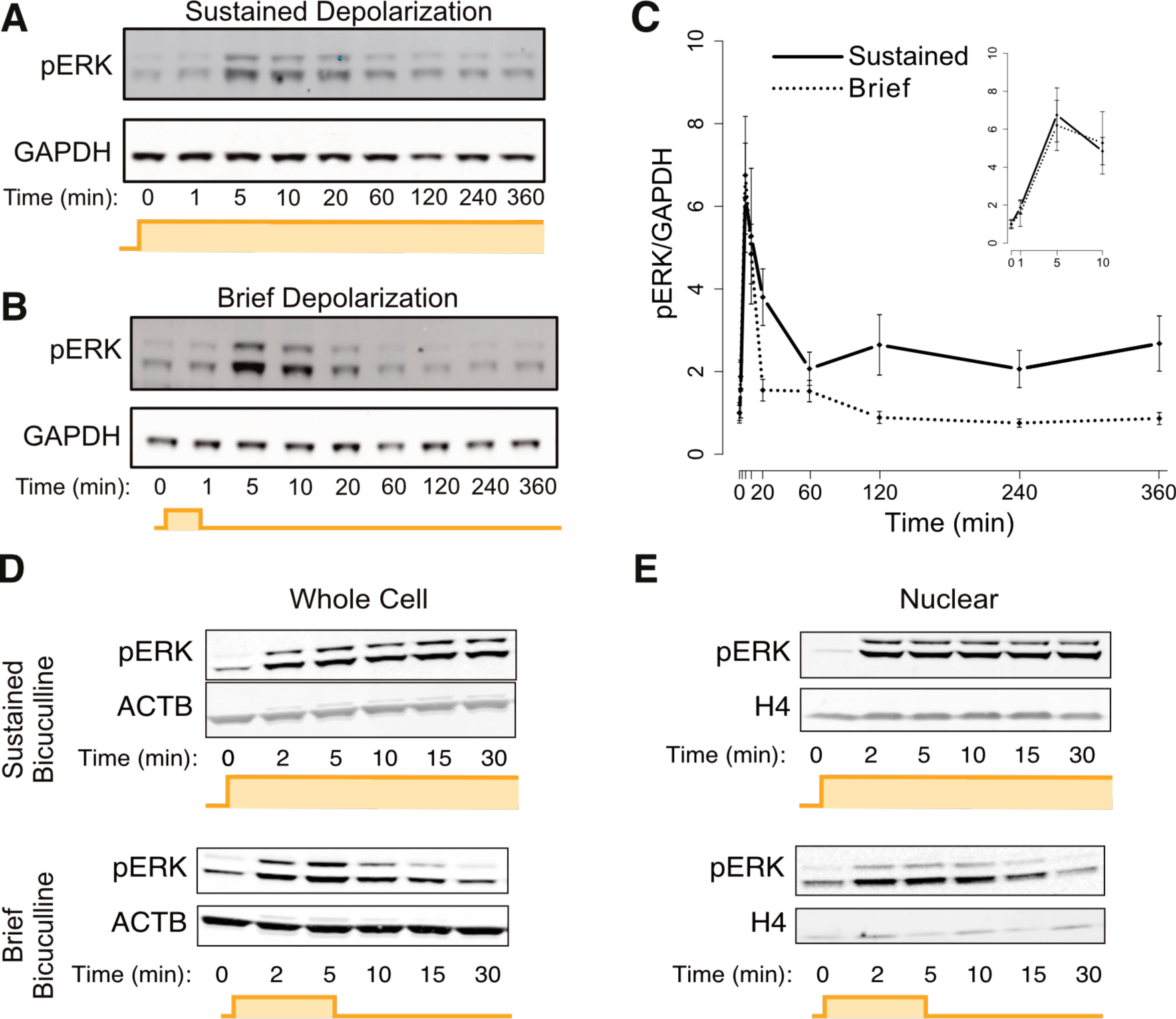
MAPK/ERK is rapidly and maximally activated by brief activity. (A) Representative western blot using an antibody recognizing phosphorylated ERK (pERK). Upper and lower bands are the phosphorylated p44 and p42 ERK paralogs (ERK1 and ERK2), respectively. Mouse cortical neurons were activated with sustained 55 mM KCl-mediated depolarization. (B) Same as in A, but with a brief (1 min) neuronal activation, followed by washout with conditioned medium. (C) Quantification of panels A-B, n=3 biological replicates. The inset is a magnified version of the first ten minutes of the same data. pERK induction at its peak (five minutes) is not statistically different between brief and sustained stimulus (p=0.3, paired two-sided *t*-test). Error bars represent +/˗ SEM. (D) Similar to A, but rat cortical neurons treated with sustained or brief bicuculline/4AP, with silencing with 2uM TTX after brief stimulation. One of n=3-4 representative biological replicates is shown. (E) Similar to D, but from isolated nuclei. One of n=3-4 representative biological replicates is shown.

We therefore hypothesized that the MAPK/ERK pathway is required for rapid PRG induction. To test this hypothesis, we measured ARG induction in the presence of MAPK/ERK pathway inhibition using the potent and highly specific allosteric MEK inhibitor, U0126 (Favata et al., 1998). We first confirmed that U0126 blocked KCl-induced ERK phosphorylation (Figure S3A). Next, we assessed ARG expression in U0126-treated neurons exposed to brief or sustained activity. Strikingly, we found that MEK inhibition dramatically blunted induction of rapid PRGs but not delayed PRGs, with 95% of rapid PRGs but only 17% of delayed PRGs sensitive to MEK inhibition (Figure 3A-C, based on > 40% decrease in maximum expression). In contrast, we confirmed that MEK inhibition had no effect on the basal, pre-stimulation ARG expression (Figure 3B) that could arise from activity-independent expression. This blunting of gene induction is unlikely to be due to off-target effects of U0126, since the MEK inhibitor PD184352 and the ERK inhibitor 11e had similar effects (Figure S3F-G). Analysis of pre-mRNA using intron reads from total RNA-seq revealed that inhibition of MAPK/ERK signaling also blunts rapid PRG expression more than delayed PRG expression at the level of transcription (Figure S3B). We also noticed that while rapid PRG mRNA induction was almost completely blocked by MEK inhibition at 20 minutes of neuronal activation, it was merely blunted at 60 minutes (Figure 3C) and barely affected at 4 or 6 hours (Figure 3C, S3D-E). Therefore, inhibition of MAPK/ERK signaling both blunts and delays the induction of first-wave genes, such that their kinetics more closely resemble those of second wave genes.

**Figure 3.**
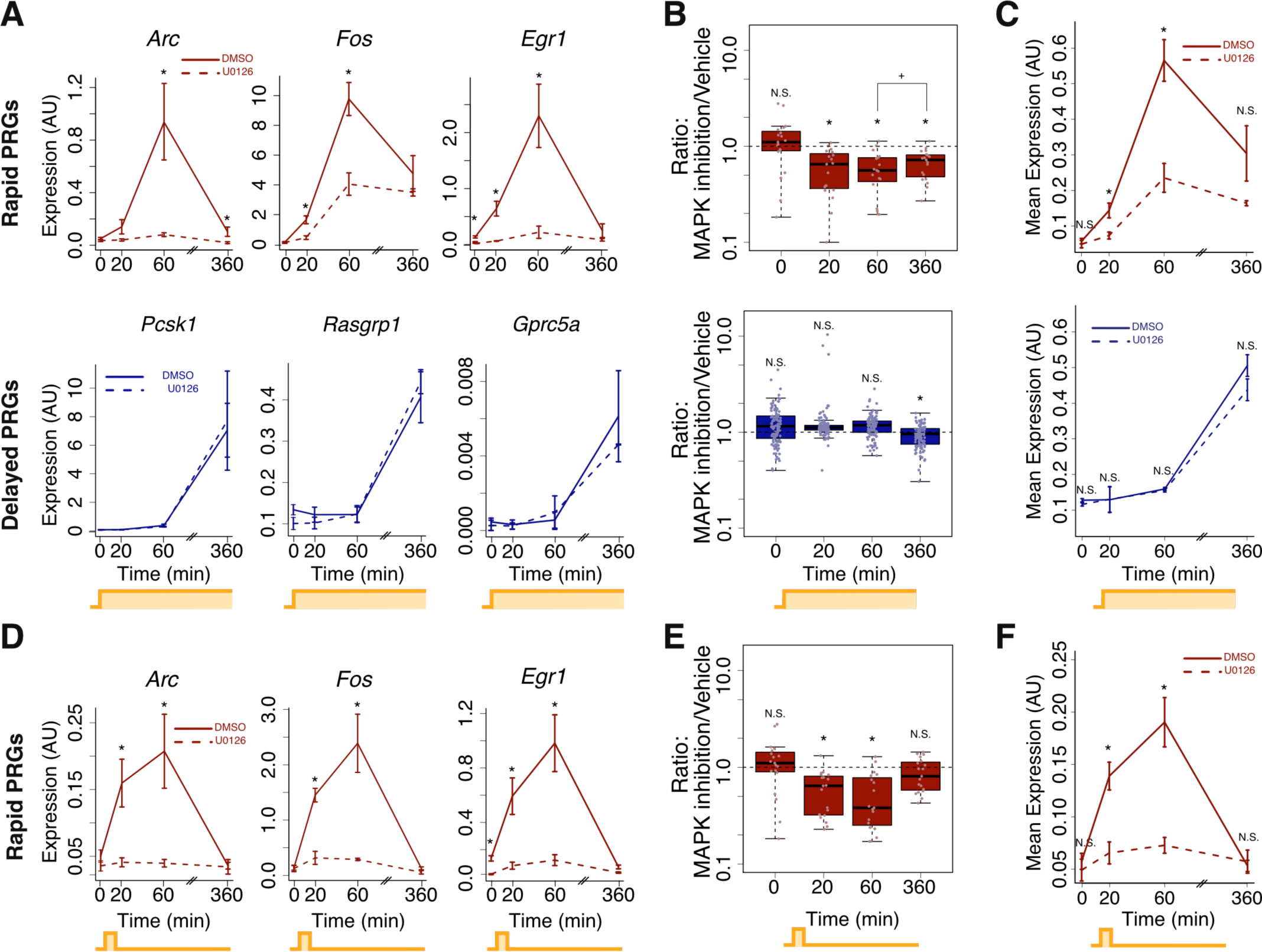
MAPK/ERK is required for the first but not subsequent waves of gene induction. (A) ARG-seq-based gene expression of three representative rapid PRGs (top) and three representative delayed PRGs (bottom) from sustained activation (KCl-depolarization) of mouse neurons with and without 10uM of the U0126 MEK inhibitor. n=3-7 biological replicates. Error bars are +/˗ S.E.M. (*p<0.01, rank-sum test). (B) ARG-seq-based gene expression for all rapid (top) and delayed (bottom) PRGs in the presence or absence of a MEK inhibitor (*significantly different from 1, p<0.001, rank-sum test; +p = 0.02, rank-sum test). Expression of rapid PRGs is more affected by MEK inhibition than expression of delayed PRGs (p = 0.002; rank-sum test on 17 rapid PRGs versus 110 delayed PRGs using the mean for each gene at its most induced time point across n=3-7 biological replicates). (C) Geometric mean of rapid PRG (top) or delayed PRG (bottom) expression with or without U0126 treatment. Same data as in panel B. Error bars are +/˗ SEM of the mean from each of n=3-7 biological replicates of geometric means of all genes in the category (*p<0.03, rank-sum test). (D) Same as A, top row, but with brief (1 min) KCl depolarization. Error bars are +/˗ S.E.M. (*p<0.01, rank-sum test). (E) Same as B, top row, but with brief (1 min) KCl depolarization (*significantly different from 1, p<0.001, rank-sum test). (F) Same as (C) but with brief (1 min) KCl depolarization (*p<0.01, rank-sum test). Related to Figure S3.

We next asked whether MAPK/ERK signaling is also required for gene induction in response to brief activity. Impressively, MEK inhibition with U0126 substantially decreased mRNA and pre-mRNA induction in response to brief activity (Figure 3D-F, S3C), blunting mRNA induction of all but one of the induced ARGs. Again, we observed similar results using the ERK inhibitor 11e (Figure S3F), suggesting that the U0126-mediated decrease in gene induction in response to brief activity is due to MAPK/ERK pathway inhibition. Thus, the MAPK/ERK pathway establishes the first wave of ARG induction, and it is required both for gene induction by brief neuronal activity and for rapid induction upon sustained neuronal activation.

### MAPK/ERK signaling is required for the first wave of gene induction in vivo

We next sought to test whether brief activity selectively induces rapid PRGs, but not other ARGs in vivo. We chose to use brief visual stimulation because it is sufficient for the formation of a long-term associative fear memory (Shi and Davis, 2001), a process that is generally transcription-dependent. We therefore exposed dark-housed mice to a brief (1 min) or sustained (up to 2.5 hr) visual stimulus and measured gene expression in the visual cortex (Figure 4A). Upon sustained stimulation, ARG-seq revealed induction of rapid PRGs, delayed PRGs, and SRGs, as expected based on these genes having been previously shown to be induced by light (Mardinly et al., 2016; Spiegel et al., 2014) (Figure 4B). These genes appeared to be specifically responding to visual stimulation rather than generalized stress, as we did not observe induction of the rapid PRG, *Fos*, in the prefrontal cortex in response to the same stimulus (Figure S4A). In contrast to sustained visual stimulation, brief visual stimulation induced rapid PRGs better than delayed PRGs (p=0.03, Fisher’s exact test, Figure 4B). Therefore, in vivo as well as in vitro, brief activity generally induces rapid PRGs but not delayed PRGs or SRGs.

**Figure 4.**
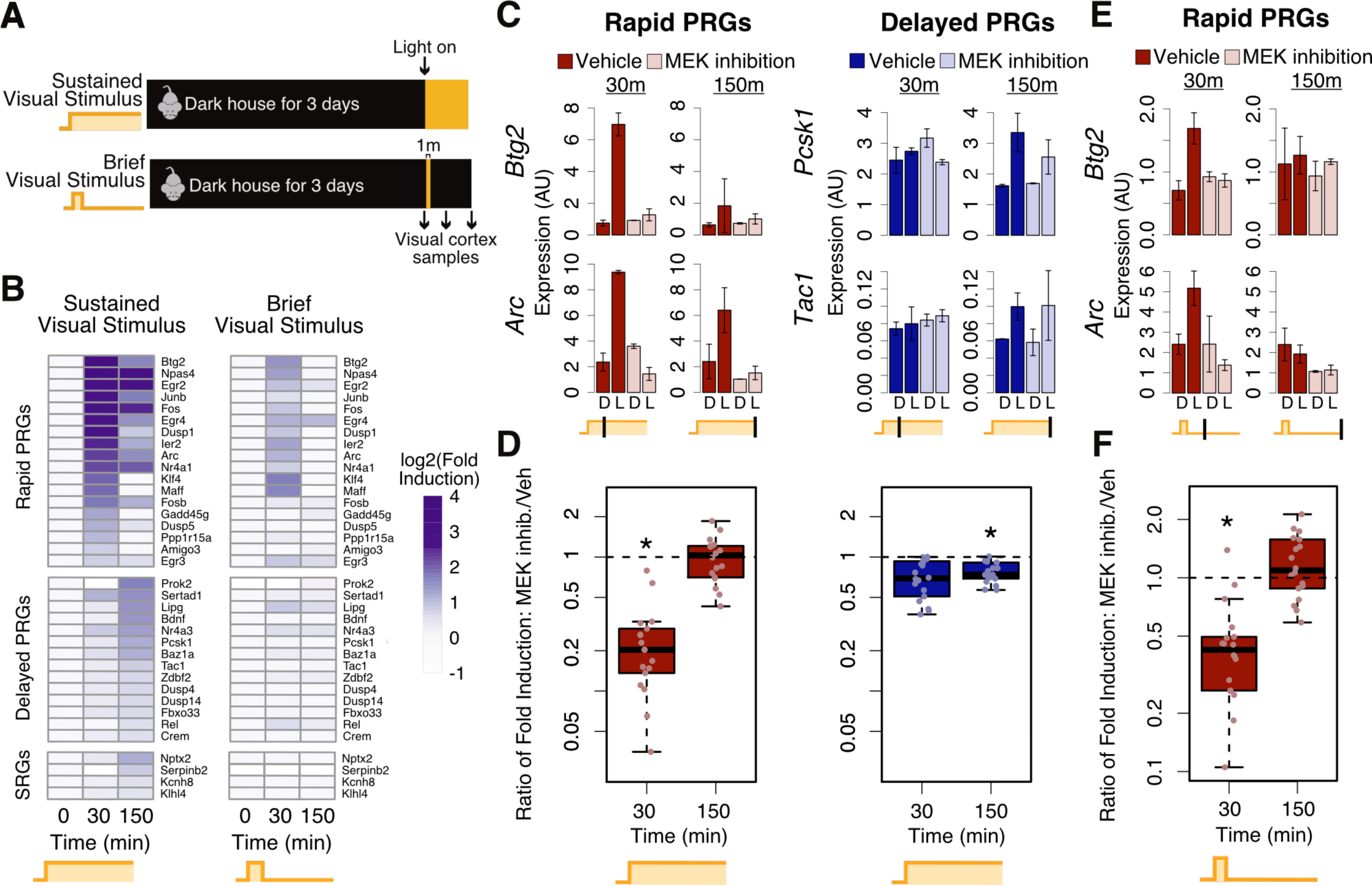
MAPK/ERK is required for the first wave but not subsequent waves of gene induction in vivo. (A) Experimental system for comparing sustained and brief visual stimulation in mouse visual cortex. Mice dark-housed for three days were exposed to sustained or brief (one minute) visual stimulation, consisting of turning on the room lights. (B) Profiling of gene expression in visual cortex before and after visual stimulation, using ARG-seq. Only genes induced >1.4 fold in any condition in vitro were included (see methods). Data are means from n=2-4 mice. Rapid PRG, delayed PRG, and SRG gene categories were defined from in vitro data as in Figure 1. Genes induced by brief visual stimulus are enriched for rapid PRGs (p=0.03, Fisher’s exact test). (C) Expression of representative rapid (left) and delayed (right) PRGs in the visual cortex in mice injected intraperitoneally with a corn oil vehicle or the MEK inhibitor SL-327 (100mg/kg), based on ARG-seq. D, dark-housed without visual stimulation. L, dark-housed with visual stimulation. Error bars are 95% confidence intervals across n = 2-3 mice. (D) Visual stimulus-mediated gene induction upon sustained stimulation with or without MEK inhibition, for all rapid PRGs and delayed PRGs detected by ARG-seq. The y-axis shows the mean KCl-dependent fold induction with MEK inhibition divided by the same fold-induction with vehicle treatment only (*i.e.*, ratio of fold-induction ratios) for each gene from n = 2-3 biological replicates (* = p<0.01 from rank-sum test assessing significance of difference from 1). Induction of rapid PRGs is more affected by MEK inhibition than induction of delayed PRGs (p = 0.02; rank-sum test, 16 rapid PRGs vs. 14 delayed PRGs using the mean for each gene at its most induced time point across n=3-5 biological replicates). (E) Expression of representative rapid PRGs in the presence of the MEK inhibitor SL-327 in mice exposed to brief visual stimulus. Details as in panel C. (F) Visual-stimulus-mediated gene induction as detected by ARG-seq upon brief stimulation with or without MEK inhibition, for all rapid PRGs, as plotted in (D) (* = p<0.01, rank-sum test based on significance of different from 1). Related to Figure S4.

We next asked whether rapid PRGs in vivo have similar chromatin characteristics as they do in vitro, as suggested by the DNaseI hypersensitivity data from 8-week cerebrum (Figure 1D). We assessed histone acetylation using ChIP-seq data from mouse hippocampus (Telese et al., 2015), since ARG induction in the hippocampus and cortex is similar (Cho et al., 2016). Compared to delayed PRG promoters, rapid PRG promoters in vivo exhibit higher levels of the active chromatin marks H4K16ac and H3K27ac, in mice not exposed to specific hippocampal activation (Figure S4B). Histone acetylation at rapid PRG promoters also occurs across a wider swath of DNA and is more bimodal in shape, suggesting lower nucleosome occupancy at rapid compared to delayed PRG promoters, as we observed in vitro. These findings suggest that rapid PRGs are primed for rapid induction by an open chromatin state in vivo.

We next investigated whether the MAPK/ERK pathway is required for rapid gene expression in vivo. We inhibited MEK by intraperitoneally injecting the blood-brain-barrier-crossing SL327, a structural analogue of U0126 (Atkins et al., 1998), into dark-housed mice 30 minutes before light exposure. Because handling or stress from the IP injection could cause gene expression on its own, we compared these mice to littermates injected with corn oil vehicle. We used western blotting to confirm that SL327 blocked ERK activation in each mouse (Figure S4C). ARG-seq of the visual cortex revealed that MAPK/ERK inhibition had a larger effect on rapid compared to delayed PRG expression when mice were exposed to sustained visual stimulation (Figure 4C-D), and it blocked nearly all ARG induction in mice exposed to brief visual stimulation (Figure 4E-F). We therefore conclude that both in vivo and in vitro, the MAPK/ERK pathway is a fast pathway necessary for ARGs to be induced rapidly or in response to brief activity.

### The MAPK/ERK pathway mediates fast Pol2 recruitment to rapid PRG promoters

Given the importance of the MAPK/ERK pathway for rapid PRG induction, we sought to understand how MAPK/ERK signaling mediates this rapid induction. We first focused specifically on the function of MAPK/ERK in regulating promoter activity. Promoters are activated in a series of steps (Adelman and Lis, 2012), each of which could be regulated by MAPK/ERK signaling. Pol2 is first recruited to promoters, followed by transcription initiation. At some promoters, transcription then pauses after a few dozen base pairs and must be released from this promoter-proximal pause into productive elongation (Adelman and Lis, 2012; Saha et al., 2011). To understand how these steps are regulated by neuronal activation, we performed Pol2 ChIP-seq in a time course following neuronal stimulation. We observed a rapid increase in Pol2 occupancy at rapid PRG promoters within 10 minutes, suggesting rapid Pol2 recruitment to these promoters (Figure 5A-B, S5A-B). At a subset of rapid PRG promoters, this recruitment occurred as soon as one minute following activation (Figure S5G). We further found that rapid PRG mRNA induction is almost completely abolished by pharmacological blockade of new initiation (Figure S5F), suggesting that initiation of newly recruited Pol2, rather than solely release of pre-bound paused Pol2, is essential for rapid PRG induction. Surprisingly, despite the slow transcriptional induction of delayed PRGs (we detect pre-mRNA only after 60m of stimulus), we observed recruitment of Pol2 to many of their promoters after 10 minutes of neuronal activation (Figure 5D-E, S5C-D), suggesting that the subsequent initiation or pause-release steps may be rate limiting for delayed PRG induction. Activity-dependent regulation of the rate of pause-release could be critically important for both rapid and delayed PRGs, but our ChIP-seq data does not enable us to detect it with confidence. These experiments indicate that the fast induction of rapid PRGs is dependent on fast recruitment and initiation of new Pol2.

**Figure 5.**
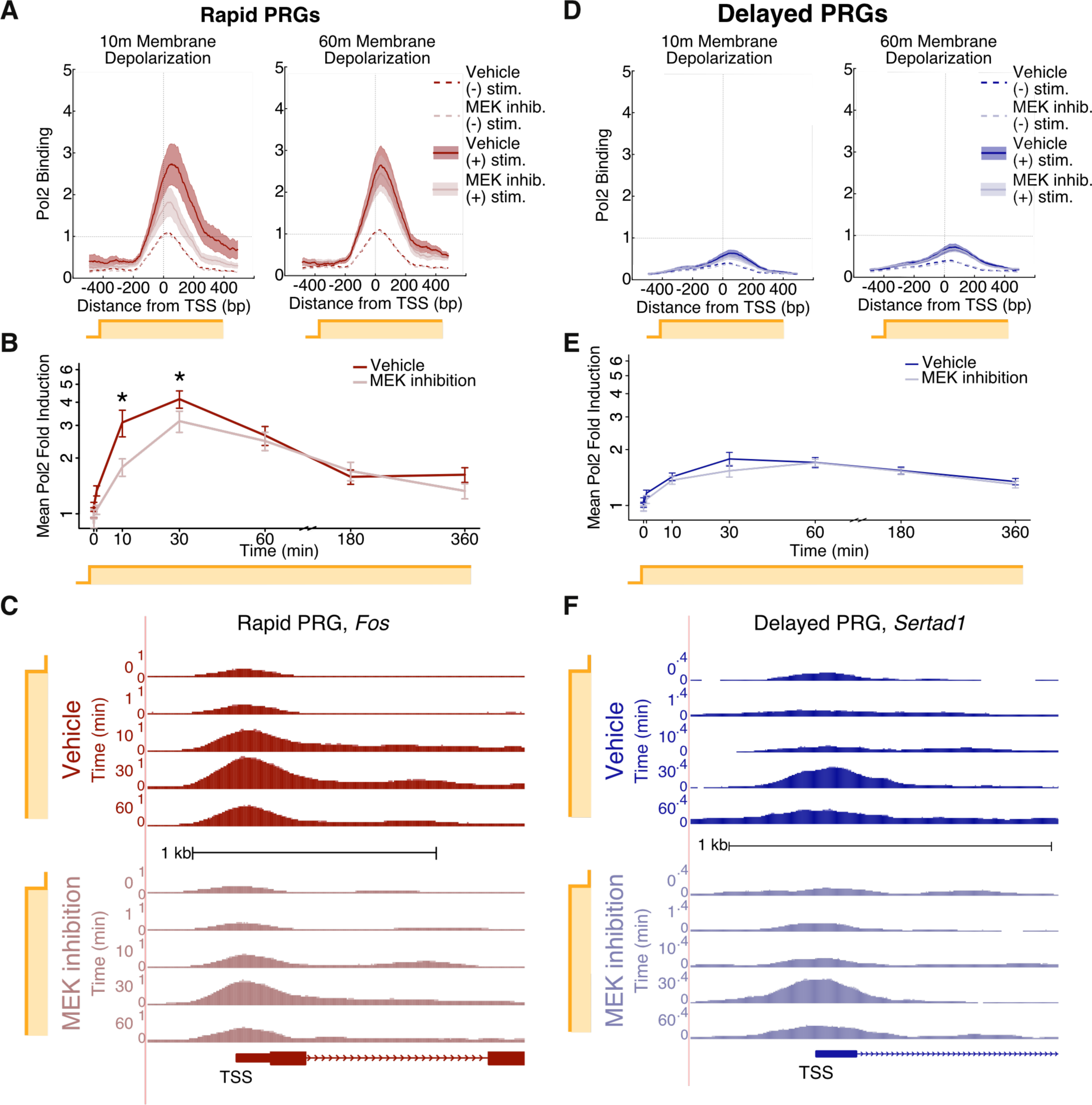
MAPK/ERK mediates rapid recruitment of Pol2 to rapid PRG promoters. (A) RNA Polymerase 2 (Pol2) binding (ChIP-seq) at the promoters of rapid PRGs, 10 and 60 minutes after KCl-mediated neuronal activation in the presence or absence of MEK inhibitor U0126 (10uM). Solid lines represent the mean and shading the S.E.M. across loci. Data shown are from one of two biological replicates. The KCl-dependent fold-increase in mean Pol2 density (˗300bp to +300bp) is significant under both vehicle and U0126 treatments (p<0.001 in each of two biological replicates, FDR-corrected paired rank sum test). (B) ChIP-seq-based time course of fold-change in Pol2 occupancy at rapid PRG promoters (˗300bp to +300bp), with or without MEK inhibition. Fold-change was calculated at each TSS using the average unstimulated Pol2 density value obtained from two DMSO- and two U0126- treated samples. Shown are mean fold-change values, with +/˗ S.E.M error bars. *FDR-adjusted p<0.01 in each of two replicates, paired rank-sum test on fold-change values. (C) Pol2 binding at the promoter of the representative rapid PRG *Fos* upon sustained neuronal activation, shown in genome browser tracks with aligned Pol2 ChIP-seq reads. The y-axis is scaled to show normalized read density from zero to the maximal value shown. (D) Pol2 binding (ChIP-seq) at the promoters of delayed PRGs, as in panel A. The KCl-dependent fold-increase in mean Pol2 density (˗300bp to +300bp) is significant under both vehicle and U0126 treatments (p<0.001 in each of two biological replicates, FDR-corrected paired rank sum test). (E) ChIP-seq-based time course of Pol2 occupancy at delayed PRG promoters. Plotting and statistics as in B. (F) Pol2 binding at the promoter of the representative delayed PRG *Sertad1* during a time course of sustained KCl treatment, based on ChIP-seq and plotted as in (C). Related to figure S5.

To assess the contribution of MAPK/ERK to the rapid recruitment of Pol2 to promoters, we performed Pol2 ChIP-seq in the presence or absence of MEK inhibition. We found that MEK inhibition does not alter Pol2 occupancy at ARG promoters in unstimulated neurons (Figure 5A-C, S5A-B), suggesting that Pol2 recruitment and pause-release at rapid PRGs is MAPK/ERK-independent in the absence of activity. In contrast, the increased Pol2 occupancy that we observed at rapid PRG promoters after ten minutes of neuronal activation is blunted by MEK inhibition (Figure 5A-C, S5A-B), suggesting that MAPK/ERK facilitates fast activity-dependent Pol2 recruitment. This dependence on MAPK/ERK for inducible Pol2 recruitment is only evident at early time points following neuronal activation (Figure 5A-C, S5A-B), suggesting that other pathways recruit Pol2 at later time points. In contrast to rapid PRGs, the relatively modest recruitment of Pol2 to delayed PRG promoters is not reproducibly affected by MEK inhibition at early or late time points for both the full set of delayed PRGs (Figure 5D-F, S5C-D) or a restricted set with greater Pol2 occupancy (FDR > 0.01, rank-sum test, see methods). These results are consistent with a model in which MAPK/ERK signaling facilitates rapid Pol2 recruitment at promoters that are primed by pre-bound transcription factors or paused Pol2 (Gilchrist et al., 2010).

### The MAPK/ERK pathway is required for eRNA transcription but not H3K27 acetylation at rapid enhancers

One mechanism by which Pol2 could be recruited to the promoters of rapid PRGs in a MAPK/ERK-dependent manner is via delivery from genomic enhancers (Szutorisz et al., 2005). We therefore asked whether enhancer activation might be dependent on MAPK/ERK signaling. We used H3K27 acetylation (H3K27ac) as a proxy for enhancer activity (Creyghton et al., 2010; Rada-Iglesias et al., 2011) and performed ChIP-seq throughout a time course of neuronal activation using an antibody against H3K27ac. We specifically focused on measuring H3K27ac levels at 940 putative ARG enhancers, defined as neuronal-transcription-factor-bound loci that are within 100kb of an ARG TSS (Kim et al., 2010). We chose this 100 kb threshold because 80% of enhancers regulate transcription start sites (TSSs) within 100kb (Chepelev et al., 2012). H3K27ac ChIP-seq revealed that 248 of the putative enhancers reproducibly gain H3K27ac within the first hour of stimulation in two biological replicates (>1.3 fold change, Figure 6A-C). Surprisingly, 70% of these 248 enhancers gain H3K27ac within the first 10 minutes of sustained activity and 85% within the first 30 minutes (Figure 6B, >1.3 mean fold change). We observed this rapid increase in H3K27ac at enhancers positioned near both rapid and delayed PRGs (Figure 6A-B). H3K27ac ChIP-seq performed in the presence of U0126 revealed that MEK inhibition had no effect on activity-dependent H3K27ac at these putative ARG enhancers, including those near MEK-dependent rapid PRGs (Figure 6D). Although MEK inhibition slightly reduces basal enhancer acetylation near rapid PRGs (Figure S6A), this relatively modest reduction is unlikely to confound our ability to assess MEK inhibition on inducible acetylation. Thus, H3K27ac is neither MAPK/ERK-dependent nor kinetically distinguishes enhancers near rapid versus delayed PRGs.

**Figure 6.**
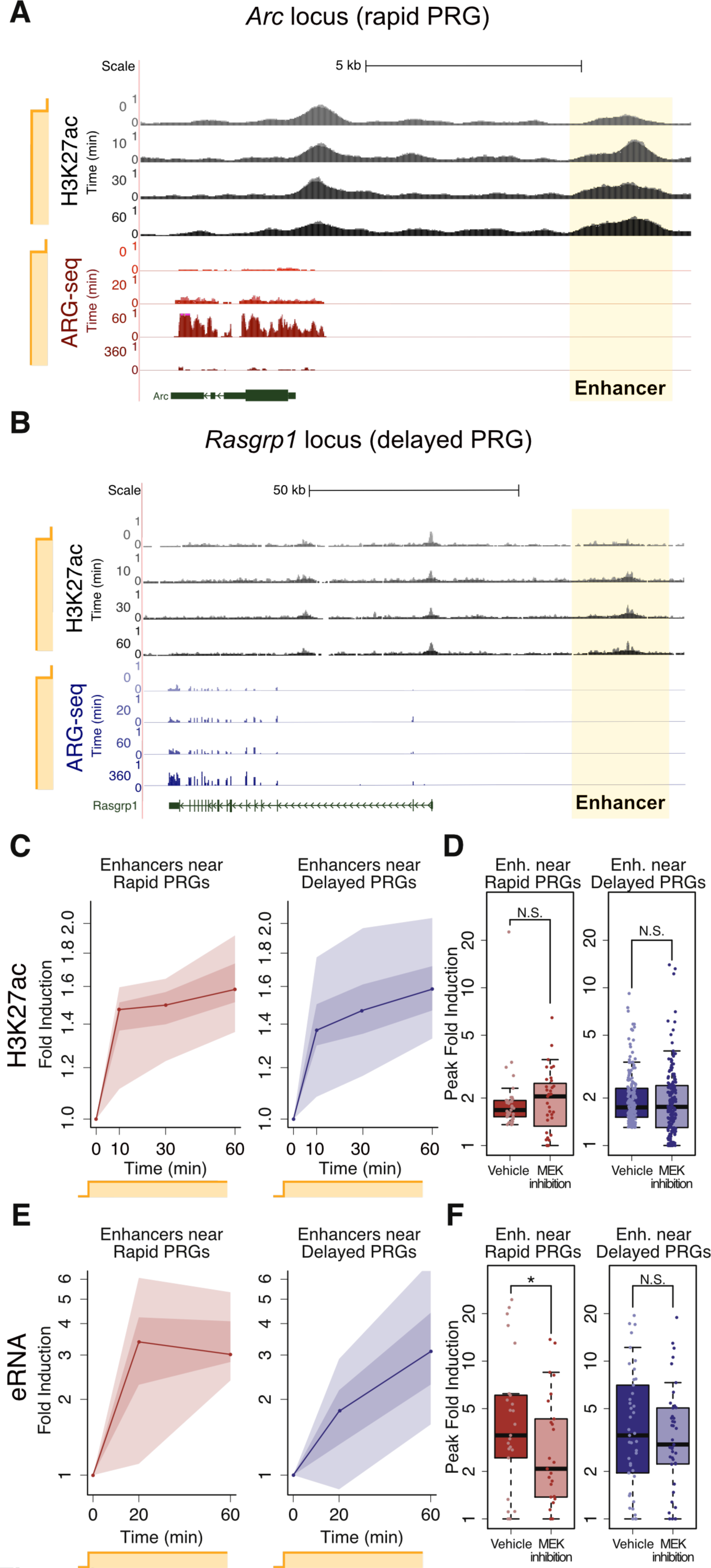
MAPK/ERK is required for rapid eRNA induction but not H3K27 acetylation at enhancers. (A) H3K27ac occupancy at the rapid PRG *Arc* locus upon sustained activation with KCl, based on H3K27ac ChIP-seq, normalized by read depth prior to visualization. The gene expression of *Arc* based on ARG-seq is shown for comparison and was also normalized across samples and re-scaled for visualization. For each sample, the duration of sustained KCl treatment is shown on the y-axis. (B) H3K27ac occupancy at the delayed PRG *Rasgrp1* locus upon sustained activation, based on H3K27ac ChIP-seq. The gene expression of *Rasgrp1* based on ARG-seq is shown for comparison. Plotting as in (A). (C) H3K27ac occupancy at enhancers near rapid PRGs and delayed PRGs upon neuronal activation with KCl, based on H3K27ac ChIP-seq. An average H3K27 acetylation change for n=2 replicates was obtained for each enhancer. These averages were plotted, with the lines representing the median, dark shading the two middle deciles, and light shading the upper and lower quartiles of values across enhancers. The increase from 0 to 10 min is significant for both enhancers near rapid PRGs and those near delayed PRGs (p<0.00001, rank-sum test using means for each enhancer from n=2 biological replicates). (D) H3K27 accumulation at enhancers near rapid and delayed PRGs is not significantly affected by MEK inhibition, as shown in box-and-whisker plots (p > 0.2, rank-sum test). The y-axis shows the mean fold induction from n=2 biological replicates at each enhancer’s most-induced time point (10, 30, or 60 min) in each condition. (E) eRNA fold induction at enhancers near rapid and delayed PRGs upon neuronal mactivation, based on eRNA-seq. Plotted as in C. (F) MEK inhibition blocks eRNA transcription at enhancers near rapid but not delayed PRGs, as shown by total RNA-seq. The y-axis shows the mean fold induction from n=2 biological replicates at each enhancer’s most-induced time point (20 or 60 min) in each condition (*, p=0.01, rank-sum test, using means for each enhancer from n=2 biological replicates; N.S., p > 0.05). Related to figure S6.

We next considered the possibility that H3K27ac may occur before or in parallel with other enhancer-activating mechanisms that might differentiate rapid and delayed PRGs. We therefore assessed another proxy of enhancer activity, enhancer RNA (eRNA) transcription (Kim et al., 2010). Surprisingly, total RNA-seq revealed that eRNA is induced more rapidly at enhancers near rapid PRGs than at enhancers near delayed PRGs, thus mirroring mRNA expression kinetics more closely than H3K27 acetylation (Figure 6E). Furthermore, in contrast to our finding that H3K27ac accumulation was not affected by MEK inhibition, MEK inhibition attenuated eRNA induction at enhancers near rapid PRGs (Figure 6F), mirroring the MAPK/ERK dependence of mRNA induction. These results indicate that rapid PRGs are distinguished by their proximity to rapid enhancers whose eRNA induction but not H3K27 acetylation is MAPK/ERK-dependent.

The finding that eRNA induction is more rapid at enhancers near rapid PRGs than at those near delayed PRGs led us to ask whether this rapidity is inherent to the enhancers or a by-product of promoter activity. To answer this question, we needed to assess enhancers individually rather than in groups based on the kinetics of nearby promoters, as we did above. However, total RNA-seq does not enable accurate classification of individual enhancers based on their eRNA expression due to the relatively small number of sequencing reads that align at each enhancer. We therefore developed an eRNA targeted capture method, eRNA-seq. eRNA-seq enriches total RNA-seq libraries for eRNAs using probes to pull down transcripts synthesized within 500 base pairs of putative ARG enhancers (Figure 7A, Table S1). Whereas in total RNA-seq, these targeted eRNAs accounted for 0.012% of aligned reads, they accounted for 6% in eRNA-seq, a 500-fold increase (Figure 7B). Of the 940 putative enhancers analyzed, 351 had quantifiable eRNA transcripts, and 89 showed a significant increase eRNA expression levels upon neuronal activation (FDR<0.1). eRNA-seq confirmed our findings from total RNA-seq: enhancers near rapid PRGs generally exhibit faster eRNA induction than enhancers near delayed PRGs (Figure S6B). These results gave us confidence that eRNA-seq provides reliable quantification of eRNA dynamics.

**Figure 7.**
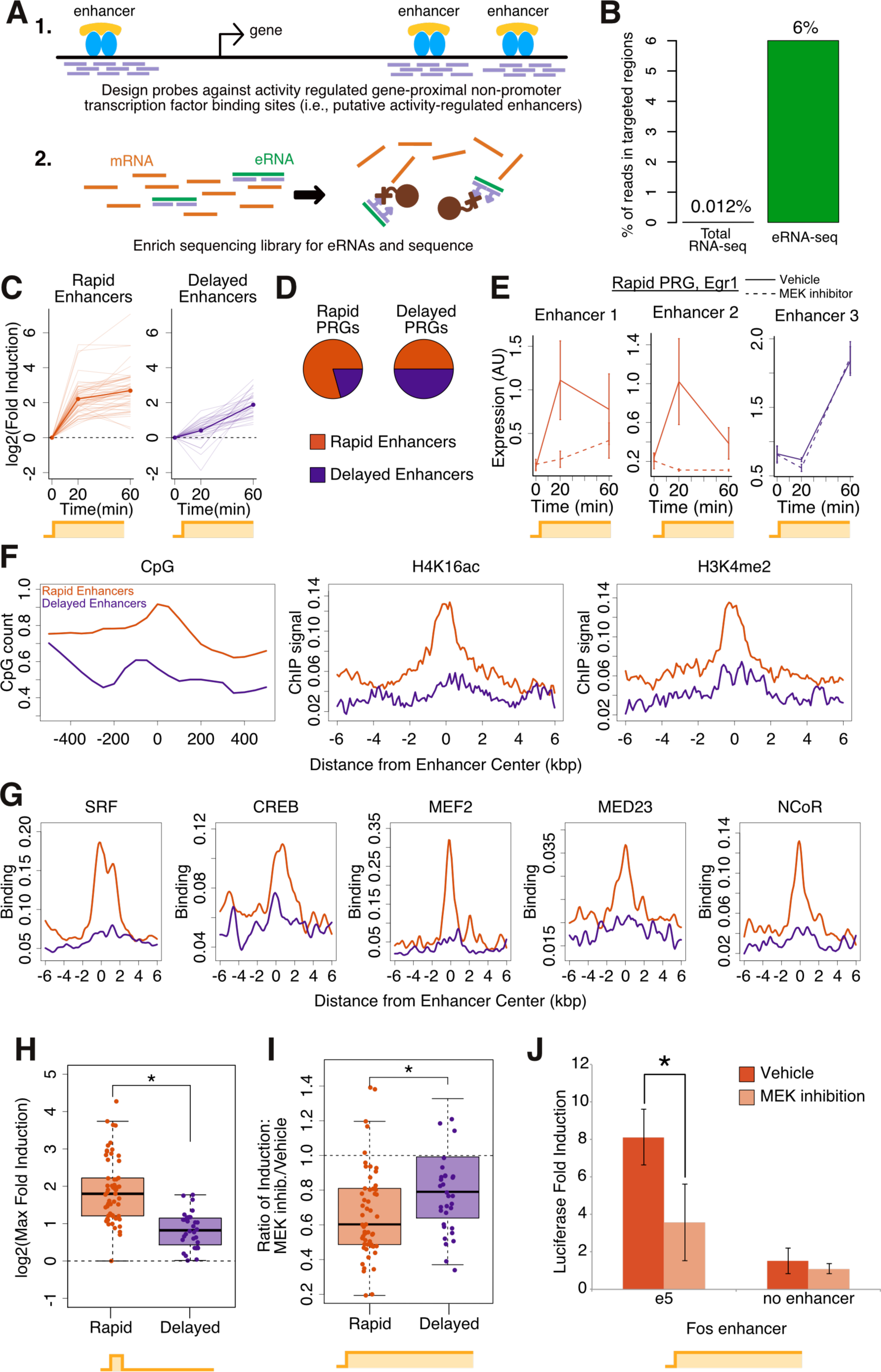
eRNA-seq enables eRNA quantification at individual enhancers, revealing rapid and delayed enhancers. (A) eRNA-seq methodology. Biotinylated RNA probes from genomic sequences within 500 bp of 940 putative activity-regulated enhancers in the mouse genome were generated via in vitro transcription from DNA oligonucleotide libraries. The probes were used to capture eRNA transcript-derived clones from total RNA-seq libraries prior to sequencing. (B) eRNA-seq increases the fraction of detectable eRNA transcripts by a factor of ~500, compared to total RNA-seq. (C) eRNA-seq-based eRNA expression at rapid and delayed enhancers upon sustained activation. Only enhancers that are significantly induced were included (FDR<0.05 at any time point). Rapid enhancers are significantly induced by 20 minutes and delayed enhancers only by 60 minutes. Light lines are means for individual enhancers from n=4 biological replicates, and heavy lines are the geometric means for all enhancers shown. (D) Rapid compared to delayed PRGs are enriched for the presence of nearby rapid enhancers (p=0.02, Fisher’s exact test), but there are also many rapid enhancers near delayed PRGs. (E) eRNA-seq-based eRNA expression at three *Egr1* enhancers, in response to sustained neuronal activation, revealing two rapid and one delayed enhancer at this rapid PRG locus. eRNA induction at enhancers 1-2 but not 3 is affected by MEK inhibition by U0126 (p < 0.05, rank-sum test, error bars are means +/˗ S.E.M.). (F) CpG content, H4K16ac, and H3K4me2 occupancy prior to stimulation at rapid versus delayed enhancers, with metaplots showing the geometric mean. CpG content, H4K16ac and H3K4me2 are significantly different between rapid PRGs and delayed PRGs or SRGs (p<10^˗7^, rank sum test using area under the curve). H4K16ac and H3K4me2 ChIP-seq data from Telese et al., 2015. (G) Binding of transcription factors, the mediator subunit MED23, and NCoR at rapid versus delayed enhancers prior to stimulation, shown as in (F). SRF, CREB, MEF2, MED23, and NCoR are significantly different between rapid PRGs and delayed PRGs (or SRGs) (p<10^˗4^, rank sum test on area under the curve). Data from Kim et al., 2010; Telese et al., 2015. (H) Rapid enhancers show greater induction in response to brief (1 min KCl) stimulus than delayed enhancers, based on eRNA-seq (p < 10^˗9^, rank-sum test). The y-axis shows the mean fold induction from n=4 biological replicates for each enhancer at its most-induced time point (20 or 60 min). (I) Rapid enhancers are more MAPK/ERK-dependent than delayed enhancers, based on eRNA-seq. For each class of enhancers, the earliest time point at which that class exhibits significant eRNA induction is shown (20 min for rapid and 60 min for delayed enhancers) (p = 0.006, rank-sum test, using means for each enhancer from n=4 biological replicates). The y-axis shows the KCl-dependent fold induction with MEK inhibition divided by the same fold-induction with vehicle treatment only (*i.e.*, ratio of fold-induction ratios). (J) Effect of MEK inhibition on the enhancer function of the *Fos* enhancer e5, using a luciferase reporter assay in which the enhancer drives transcription from a minimal *Fos* promoter. * p<0.03 from *t*-test based on n=3 replicates. Related to figure S6.

We next applied eRNA-seq to assess eRNA dynamics at individual enhancers. We defined rapid enhancers as those with significant eRNA induction at 20 minutes of sustained activity and delayed enhancers as those with significant induction only at 60 minutes (FDR<0.1, Figure 7C). We found that 79% of enhancers near rapid PRGs are rapid enhancers (Figure 7D-E), consistent with the idea that rapid enhancers may contribute to rapid PRG induction. However, 21% of enhancers near rapid PRGs exhibit delayed kinetics, consistent with an observation in Drosophila embryos that enhancers regulating a single gene may independently control distinct phases of promoter activation (El-Sherif and Levine, 2016). Surprisingly, half of the activity-regulated enhancers near delayed PRGs are rapid enhancers (FFigure 7D), again supporting the idea that enhancers are regulated independently of their nearby promoter. Consistent with a possible role for rapid enhancers in regulating the rapidity of rapid PRG induction, we found that rapid enhancers exhibit greater induction in response to brief activity than delayed enhancers (Figure 7H). This greater induction of rapid enhancers by brief neuronal activation was observed even when comparing only the rapid and delayed enhancers that are located near delayed PRG loci (p<10^˗4^, rank-sum test, see methods). This dissociation of enhancer kinetics with mRNA kinetics suggests that the rapid induction and sensitivity to brief stimulus characteristic of rapid enhancers is intrinsic to the enhancers themselves.

Having found that enhancers show enhancer-intrinsic induction kinetics, we asked how rapid enhancers are activated more quickly than delayed enhancers. Compared to delayed enhancers, we found that rapid enhancers have significantly higher CpG content, higher levels of active chromatin marks, greater DNAse hypersensitivity, and greater binding of the transcription regulators SRF, MEF2 and Mediator in unstimulated neurons (Figure 7F-G, S6C-D). In addition, rapid but not delayed enhancers bind the transcriptional repressor NCoR in unstimulated neurons (Figure 7G, S6F), suggesting active repression. Consistent with the suggestion of active repression by NCoR, rapid and delayed enhancers in unstimulated neurons have similar levels of eRNA expression (Figure S6E) despite the more active chromatin state at rapid enhancers. The more active chromatin state at rapid enhancers appears to be intrinsic to the enhancers rather than an indirect effect of their associated promoters, since a comparison of just those rapid and delayed enhancers near delayed PRGs revealed the same differences in CpG content, active chromatin marks and transcription factor pre-binding in unstimulated neurons (p<0.01, rank-sum test, see methods). Despite these differences between rapid and delayed enhancers in unstimulated neurons, after two hours of neuronal activation, rapid and delayed enhancers showed similar increases in Pol2 occupancy (Figure S6C), consistent with our observation that both classes of enhancers are transcribed at later time points. Using eRNA-seq in the presence of a MEK inhibitor, we also found that rapid enhancers are more MAPK/ERK dependent than delayed enhancers (Figure 7E,I). These results indicate that rapid enhancers are primed for rapid MAPK/ERK-dependent activation whether they are near first- or second-wave genes.

Given our finding that rapid enhancers require MAPK/ERK signaling for eRNA induction, we asked whether rapid enhancers also require MAPK/ERK signaling to enhance promoter transcription. We paired a previously characterized *Fos* enhancer (“e5”) (Joo et al., 2015) with a minimal *Fos* promoter driving luciferase gene expression. As expected, the enhancer was able to enhance promoter activity in an activity-dependent manner (Figure 7J). Interestingly, MEK inhibition blocked this e5-dependent induction of promoter activity. These results suggest that for a subset of enhancers, MAPK/ERK signaling is required to drive mRNA expression from their target promoters.

## DISCUSSION

Using genome-scale technology, we find that a neuron’s activity pattern history is encoded in its gene expression profile. Sustained neuronal activity induces three temporal waves of gene induction: rapid PRGs, delayed PRG, and SRGs. In contrast, brief activity induces only the first of these waves, i.e., rapid PRGs. Rapid PRGs are distinguished from delayed PRGs and SRGs by a constitutively open chromatin state at their promoters and pre-binding of transcription factors and Pol2. Rapid PRGs are also uniquely dependent on MAPK/ERK signaling for their induction (Figure 8). MAPK/ERK signaling can activate rapid PRG promoters via activation of rapid enhancers. eRNA-seq reveals that the fast eRNA induction at these rapid enhancers distinguishes them from delayed enhancers, similar to the distinction between rapid and delayed PRGs. Surprisingly, at rapid enhancers, MAPK/ERK signaling is required for eRNA but not H3K27ac induction. MAPK also mediates the fast recruitment of Pol2 to rapid PRG promoters. Abolishing MAPK/ERK signaling not only alters the multi-wave structure of the ARG response by blunting and delaying rapid PRG induction, but it also abolishes rapid PRG induction in response to brief activity. In this way, MAPK/ERK both establishes the multi-wave structure of ARG transcription and enables activity-pattern-specific gene induction. This shared mechanism suggests that a biological advantage of the multi-wave structure of activity-regulated gene expression is to enable different activity patterns to induce distinct subsets of genes.

**Figure 8.**
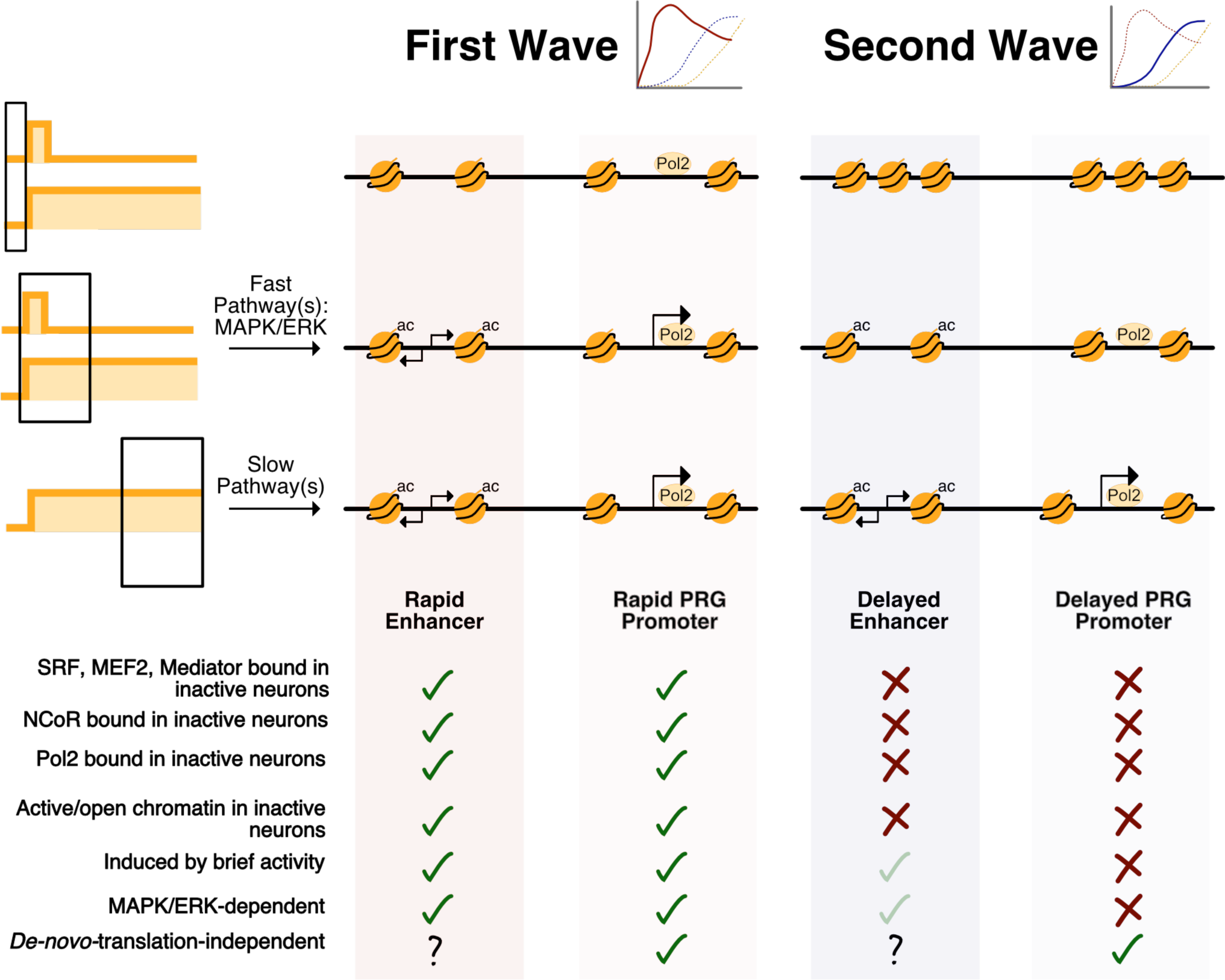
Distinguishing features of first wave genes (rapid PRGs) and second wave genes (delayed PRGs). Rapid PRGs are distinguished by an open chromatin state, proximity to rapid enhancers, and dependence on MAPK/ERK signaling. MAPK/ERK facilitates rapid Pol2 recruitment to rapid PRG promoters but is not required for Pol2 recruitment to delayed PRG promoters. Rapid eRNA induction occurs at rapid enhancers in a MAPK/ERK-dependent manner, but subsequent eRNA induction at delayed enhancers is less MAPK/ERK-dependent. Light green check marks indicate partial effects.

### MAPK/ERK establishes the first wave of gene induction

Our results indicate that the MAPK/ERK pathway is a key structural determinant of the first wave of gene induction. Our identification of MAPK/ERK as a fast pathway for gene induction could inform studies of the many human neurodevelopmental disorders caused by mutations in genes that are part of or interact with this pathway (Jindal et al., 2015; Thomas and Huganir, 2004). We show that MAPK/ERK signaling is most important for rapid PRG transcription and Pol2 recruitment at early time points and in response to brief activity. This finding is in contrast to previous studies that suggested the MAPK/ERK pathway could be a relatively slow regulator of transcription that does not respond to brief activity (Hardingham et al., 2001; Murphy et al., 1994; Toettcher et al., 2013; Wu et al., 2000). MAPK/ERK is slow to phosphorylate the transcription factor CREB in studies that detected CREB phosphorylation in bulk using immunocytochemistry (Wu et al., 2000) or western blotting (Hardingham et al., 2001). However, it is possible that CREB phosphorylation may occur quickly at rapid PRGs but be detectable in bulk only later, when it occurs more broadly. Alternatively, MAPK/ERK may instead activate rapid PRGs via SRF (Ramanan et al., 2005) and Elk-1 (Xia et al., 1996). Despite its slow phosphorylation of CREB, others have found that the MAPK/ERK pathway can be rapidly activated in the nucleus in response to transient stimulation (Dudek and Fields, 2001; Zhai et al., 2013) and is required for induction of several genes that we can now classify as rapid PRGs (Davis et al., 2000; Eriksson et al., 2006; Zheng et al., 2009). MAPK/ERK signaling is also required for late-phase LTP, a transcription-dependent process that occurs in response to brief activity (Huang et al., 2000). Finally, the MAPK/ERK pathway architecture is well suited to inducing the first wave of transcription: the multiple kinases as well as the potential for positive feedback in the pathway (Kholodenko et al., 2010) are ideal for amplifying transient signals. This ability could explain both the rapidity of the MAPK/ERK response and its responsiveness to brief activity and may be the foundation for its ability to store a memory of neuronal activity for long enough to activate transcription.

There are at least two non-mutually-exclusive ways that MAPK/ERK could specify which genes are included in the first wave. In a passive model, rapid PRG promoters could be uniquely sensitive to MAPK/ERK signaling solely due to their open chromatin state. This model is similar to the observation in macrophages that the a subset of PRGs have open chromatin and prebound Pol2 at their promoters, obviating the need for chromatin remodeling, which is required at other PRGs and SRGs (Hargreaves et al., 2009; Ramirez-Carrozzi et al., 2009, 2006). Because MAPK/ERK signaling is most active in early stages of the response to neuronal activity, it may selectively act only on promoters and enhancers that have open chromatin at these early time points. In an active model, MAPK/ERK signaling could activate rapid PRGs due to specific binding of MAPK/ERK-dependent transcriptional activators at these genes. Indeed, we observed that rapid PRG promoters, but not delayed PRG or SRG promoters, bind the MAPK/ERK-regulated transcription factor SRF (Treisman, 1996). SRF is required in vivo for the transcription of genes we classify here as rapid PRGs (Ramanan et al., 2005). SRF often acts in concert with another transcription factor, Elk-1, which is directly phosphorylated by MAPK/ERK (Gualdrini et al., 2016; Marais et al., 1993; Sgambato et al., 1998; Sharrocks, 1995; Xia et al., 1996). Elk-1 has been shown to facilitate Pol2 recruitment via interactions with the same Mediator subunit MED23 (Allen and Taatjes, 2015; Besnard et al., 2011; Wang et al., 2005) that we found selectively pre-bound to rapid PRG promoters and rapid enhancers. A third function for MAPK/ERK signaling that could apply to either model is phosphorylation of histone tails either directly, via its downstream kinase MSK (Josefowicz et al., 2016), or via interactions with Elk-1 (Esnault et al., 2017).

### Separable mechanisms of enhancer activation revealed by MAPK/ERK

Surprisingly, we find that the MAPK/ERK pathway regulates eRNA induction but not H3K27ac accumulation at rapid enhancers, suggesting that enhancer activation occurs in multiple mechanistically separable steps. H3K27ac is a commonly used mark for enhancer activity (Creyghton et al., 2010), but we find H3K27ac accumulates at enhancers even in the presence of MAPK/ERK inhibition, which blocks eRNA (and mRNA) induction. Enhancer activation may begin with rapid acetylation and be followed by MAPK/ERK-dependent eRNA transcription and promoter activation. Alternatively, acetylation and eRNA induction might occur independently and contribute to distinct mechanisms by which enhancers activate promoters. Either of these models appear discrepant with a recent study suggesting that eRNA transcription is required for acetylation at non-inducible gene loci (Bose et al., 2017). This apparent discrepancy could be a consequence of eRNA being important for H3K27ac maintenance rather than initiation. Indeed, in other contexts histone acetylation has been shown to accumulate despite blocking eRNA transcription, Pol2 recruitment, or initiation of transcription (Hah et al., 2013; Kaikkonen et al., 2013; Wang et al., 2005). These, as well as other analyses (Zhu et al., 2013) also suggest that eRNA transcription may be a better marker for enhancer activation than H3K27ac, more accurately reflecting an the extent to which an enhancer is activating transcription at a nearby promoter. Given these findings, eRNA-seq may be a particularly useful technique for reliably assaying enhancer activation genome wide, as it was previously impractical, laborious, or costly to quantify eRNA expression at many enhancers using total RNA-seq, GRO-seq, or NET-seq (Churchman and Weissman, 2011; Core et al., 2008).

### Regulation of delayed PRGs and SRGs

While the MAPK/ERK pathway establishes the first wave of gene expression, our results suggest that other pathways must establish later waves of gene induction. This idea is in contrast to the dominant role of the MAPK/ERK pathway in PC12 cells, where differences in MAPK activity are sufficient to drive distinct cellular outcomes: brief activation causes proliferation while sustained activation causes differentiation (Gotoh et al., 1990; Marshall, 1995; Santos et al., 2007). In contrast, our results suggest that in neurons the cell biological effects of the later waves of transcription require a different cell-signaling pathway that is activated in response to sustained activity. It is possible that this pathway is the CaMKIV pathway given that it is required for synaptic scaling, which only occurs response to sustained neuronal activity (Ibata et al., 2008). A late-wave-regulating pathway could be important for histone turnover (Maze et al., 2015) or chromatin remodeling at the promoters and enhancers of late-wave genes. A role for chromatin remodeling is supported by the observation that in macrophages, PRGs with closed promoters, unlike those with open promoters, require SWI/SNF remodelers for their induction (Hargreaves et al., 2009; Ramirez-Carrozzi et al., 2009, 2006). Chromatin opening is also a key step in enhancer activation not only during neuronal development (Frank et al., 2015) but also in response to neuronal activity (Su et al., 2017). Our finding that delayed enhancers have relatively inactive chromatin prior to neuronal activation makes them similar to “latent” macrophage enhancers that are slowly activated and are not pre-bound by transcription factors (Ostuni et al., 2013). Chromatin remodeling at delayed enhancers and the promoters of delayed PRGs and SRGs could be dependent on the accumulation of acetylation at these regions upon neuronal activation, either via recruitment of histone acetytransferases (Kim et al., 2010) or removal of histone deacetylases (Guan et al., 2009). This possibility is supported by the finding in neurons and in macrophages that the induction of slowly induced genes is selectively dependent on bromodomain-containing proteins that both recognize histone acetylation (Nicodeme et al., 2010; Sullivan et al., 2015) and are important for learning (Korb et al., 2015). Despite this function for histone acetylation in delayed gene induction, our results suggest that histone acetylation itself is a fast process at enhancers. However, we also find that histone acetylation is MAPK/ERK-independent, and it could instead be mediated by rapid CREB phosphorylation driven by a CaMK pathway (Wu et al., 2000). In this model, late-wave gene induction would be delayed not due to the speed of the signaling pathway that regulates it, but rather due to the energy-intensive, time-consuming requirement for chromatin remodeling.

### Role of rapid PRG protein products

The protein products of rapid PRGs are likely to be required for the cell biological changes that occur following a single occurrence of brief neuronal activity. In one potential example, the maintenance of LTP, a MAPK/ERK-dependent process (Huang et al., 2000), requires transcription within a brief critical window immediately following stimulation (Saha and Dudek, 2013), during which our data suggest that only rapid PRGs are induced. In another example, multiple electrode array (MEA) recordings of cultured hippocampal neurons have shown that just seven minutes of bicuculline treatment (similar to our five-minute treatment) is sufficient to induce long-lasting synchronous bursting, and this synchronization of bursting is dependent on both MAPK/ERK and transcription (Arnold et al., 2005). The 19 rapid PRG protein products classified here are likely candidates for mediating the synaptic changes that underlie both bicuculline-induced synchronous bursting and LTP. However, the notion that rapid PRGs are sufficient to induce changes in synaptic plasticity is somewhat puzzling given that the majority of rapid PRGs are transcription factors. On possibility is that these transcription factors prime the neuronal genome, making it easier to induce late-wave gene induction in response to subsequent brief activity. However, we hypothesize that the immediate cell biological effect of this gene program in response to brief activity is due in large part to the effect of just a few genes that are not transcription factors, such as *Arc*, which is required for LTP (Plath et al., 2006), and *Amigo3*. Our results therefore suggest that *Arc* plays a large role in the transcription-and activity-dependent changes in synaptic plasticity underling memory consolidation, and future studies might benefit from furthering understanding of this role rather than searching for new roles for other ARGs. We also identified genes that require sustained activity for their transcription, and these delayed PRGs may be key regulators of the homeostatic plasticity that occurs selectively in response to sustained activity.

## Author Contributions

K.M.T. and J.M.G designed the part of the study using KCl/visual stimulation. R.N.S. and S.M.D. separately designed the bicuculline experiments. K.M.T. performed and analyzed all RNA-seq experiments, H3K27ac ChIP-seq, western blotting for KCl experiments, high-throughput qPCR, and enhancer RT-qPCR. K.M.T. analyzed previously published genomic data. R.N.S., P.R., and M.K.W. performed the bicuculline experiments including Nanostring and western blots of nuclear extracts. J.M.W. with K.M.T. analyzed the Nanostring data. N.D.S. performed and analyzed RNA polymerase ChIP-seq and triptolide experiments. J.-H.C. performed visual stimulation experiments. S.M.C. performed 11e experiments. R.D.J. performed the luciferase assay. K.M.T. and J.M.G. wrote the manuscript with input from R.N.S., S.M.D. and N.D.S.

## Acknowledgements

We thank Karen Adelman, Charles Danko, Sandeep R. Datta, Steven Flavell, Harrison Gabel, Michael Greenberg, Elizabeth Hong, Kelly Biette, Heather Landry, Colin Waters, Emily Low, Ian Hill, and Patricia Rohs for a critical reading of the manuscript. We thank the Gray laboratory members for helpful discussion throughout the course of the project. We thank Joseph Ling and Matthew Friese for early work on this project. We also thank the BCH IDDRC, 1U54HD090255 for Fluidigm qPCR. This work is funded by R01 MH101528-01, the Kaneb family (J.M.G lab), and the Intramural Program of the National Institute of Environmental Health Sciences, National Institutes of Health Z01 ES100221 (S.M.D. lab). K.M.T. was funded by the National Science Foundation GRFP. N.R.D. was funded by the National Institute of Health NRSA. R.N.S. was funded by R00 MH096941. J.H.C. is funded by a postdoctoral fellowship from the Canadian Institute of Health Research.

## METHODS

### CONTACT FOR REAGENT AND RESOURCE SHARING

Further information and requests for resources and reagents should be directed to and will be fulfilled by the Lead Contact, Jesse Gray.

### EXPERIMENTAL MODEL AND SUBJECT DETAILS

#### Mouse primary neuronal cultures and stimulation

Cortical neurons were dissected from embryonic day 16 (E16) CD1 embryos of mixed gender. They were dissociated with papain (Worthington, (L)(S)003126) and plated on plates coated for at least one hour with poly-ornithine (30mg/mL, Sigma) in water and then washed three times with water. They were maintained in neurobasal media (Invitrogen) supplemented with B27 (Invitrogen), Glutamax (Invitrogen), and penicillin/streptomycin (Invitrogen). At 6 or 7 days in vitro (DIV) neurons were silenced with APV (100uM, Tocris) and NBQX (10uM, Tocris). 14-16 hours later neurons were stimulated with a final concentration of 55mM potassium chloride using KCl depolarization solution (170mM KCl, 10mM Hepes pH 7.4, 1mM MgCl_2_, 2mM CaCl_2_). For sustained stimulation, KCl was left on neurons for up to 6 hours, whereas for brief stimulation, it was added for one minute, and then removed and replaced with conditioned neurobasal supplemented with APV and NBQX until RNA collection. 10μM U0126 (Tocris), 625nM 11e (Tocris), 30μMcycloheximide (Cell Signaling) or DMSO (equal volume) were added 30 minutes before stimulation and left on the neurons throughout the experiment.10μM Triptolide (Tocris) was added 5 minutes before stimulation.

#### Rat cell culture and stimulation

Cultures of cortical neurons were prepared from embryonic day 18 Sprague Dawley rats of mixed gender (NIEHS Animal Study Proposal #01-21). Dissociated cortical neurons were plated in Neurobasal medium (Invitrogen) supplemented with 25 mM glutamate (Sigma-Aldrich) and 0.5 mM L-glutamine (Sigma-Aldrich) and either B27 (Invitrogen) or NS21 and maintained in a similar medium without the glutamate. NS21 was prepared in the laboratory as previously described (Chen et al., 2008). Neurons were used routinely between 10–14 days *in vitro*. To induce gene transcription under basal conditions using synaptic circuits, we triggered neuronal activity by co-treating neurons with 50μM Bicuculline (Sigma-Aldrich) and 7.5μM 4-Aminopyridine (Acros Organics) (Papadia et al., 2005). Activity was ceased at the desired time point using 2μM TTX.

#### Mice

For *in vivo* experiments, male P70-P90 C57/BL6 mice were used.

## METHOD DETAILS

### Visual stimulation

Adult mice were housed in the dark for three days. On the day of the experiment, they were intraperitoneally injected with 100mg/kg of SL327 (Tocris) in corn oil or with a corn oil vehicle. Injections started 30 minutes before visual stimulus and continued once per hour for the duration of the experiment to maintain the effects of the drug. SL327 was solubilized first in 100% ethanol, then this was added to corn oil and vortexed for 30 minutes. The ethanol was then removed from the mixture using a speed vac. For the visual stimulation, the overhead room lights were turned on for one minute and then turned off (brief stimulus) or were turned on continuously for up to 2.5 hours. Mice were sacrificed before the stimulus or either 30 minutes or 2.5 hours after turning on the lights. Their visual cortices were immediately dissected. One hemisphere from each mouse was homogenized in Trizol (Invitrogen) for subsequent ARG-seq, and the other was homogenized in cold lysis buffer (see Western Blotting) for western blotting to confirm ERK activation.

### RNA extraction from rat neurons and qPCR

Total RNA was isolated from dissociated neurons using the RNeasy Mini Kit (Qiagen) with in-column DNase (Qiagen) digestion. cDNA was synthesized using MuLV reverse transcriptase (Promega), random primers (Promega), oligo dT primers (Promega), and RNase inhibitors (Thermo Scientific). qPCR was performed using iTaq Universal Sybr Green Supermix (BioRad) and the BIO-RAD CFX Connect realtime PCR Detection System. Pre-mRNA was estimated as described (Saha et al., 2011). Rat PCR primers used in this study have been previously described (Saha et al., 2011).

### RNA extraction from mouse neurons and qPCR

Neurons were collected in Trizol (Invitrogen), and total RNA was extracted using the RNeasy kit (Qiagen) with in-column DNase treatment (Qiagen) according to the instructions of the manufacturer. The RNA was then either used for RNA sequencing (see below) or converted to cDNA using the High Capacity cDNA Reverse Transcription kit (Applied Biosystems). For standard qPCR experiments, we used SsoFast Evagreen supermix (BioRad) with the following primers: *Fos*: GGCTCTCCTGTCAACACACA, TGTCACCGTGGGGATAAAGT, *Egr1*: GGGATAACTCGTCTCCACCA, CCTATGAGCACCTGACCACA, *Pcsk1*: TGCAGGTGAAATTGCCATGC, GGCCAGGGTTGAATCCAATTG; *Rasgrp1*: TGACAACTGTGCTGGCTTTC, TGCACTGTTTGTGGCAGTTC, *Gapdh*: CGTCCCGTAGACAAAATGGT, TCGTTGATGGCAACAATCTC or eRNA primers as in (Joo et al., 2015). For high-throughput qPCR, we used Taq-man qPCR probes (designed by Invitrogen) using the Fluidigm microfluidics system.

### NanoString

NanoString probes were designed for indicated pre-mRNAs by NanoString technologies and assays were performed following the manufacturer’s protocol.

### RNA sequencing

Before library preparation, for capture experiments, ERCC spike-in RNA (Ambion) was added to RNA samples according to the instruction of the manufacturer. Libraries were prepared using the High Throughput Total RNA TruSeq kit (Illumina), following the instructions of the manufacturer but scaling down all volumes to 1/3 of the recommended volumes. Libraries were sequenced on a NextSeq (Ilumina) to a depth of at least 30 million reads per library for total RNA-seq. We aligned reads to the mm9 genome using the STAR aligner (Dobin et al., 2013), and then made the resulting SAM files into BED files using SAMtools and BEDtools (Li et al., 2009; Quinlan and Hall, 2010). We used UCSC-tools (Kuhn et al., 2013) to make bigWig files for viewing on the genome browser. We used bedtools map to count reads in both exons and introns. We then analyzed the raw count data using R, including edgeR (Robinson et al., 2009).

### Capture RNA sequencing

For ARG-seq, capture probes were designed as oligonucleotides tiling activity-regulated exons and control exons. Genes to be captured were 229 ARGs that showed a reproducible 3.5 fold increase in transcription at either 1 or 6 hours of KCl treatment in two replicates of published RNA-seq data (Kim et al., 2010) and 47 genes that showed no change with KCl but spanned a range of expression values (controls). Synthesized probes were 100 base pairs in length, with each probe overlapping the previous probe by 76 base pairs. Probes had PCR primer binding sites and IVT promoters added. These oligonucleotides were ordered from Custom Array, PCR-amplified, and transcribed in vitro into biotinylated RNA baits using the Megascript SP6 In Vitro Transcription kit (ThermoFisher).

For eRNA-seq, capture probes were designed as oligonucleotides tiling putative activity-regulated enhancers, which were identified based on their location relative to ARGs and their transcription factor binding. To identify these putative enhancers, we started with all CREB,SRF, CBP, Npas4 or Pol2 binding sites from a previous study (Kim et al., 2010). We then took only those sites that were within 100kb of a transcription start site of one of the ARGs used in our ARG-seq experiment. We eliminated intragenic enhancers and those located within 1kb from the transcription end site or 500bp from the transcription start site of a gene. We designed probes to span the entire TF-bound putative enhancer, plus 500 bp on each side. This oligonucleotide library was ordered from Twist Biosciences. We amplified and in vitro transcribed the RNA baits as described above for the ARG-seq baits.

For ARG-seq and eRNA-seq, samples were treated in the same manner as with total RNA-seq, except that after library preparation, 250ng of pooled libraries were heated to 95C to denature DNA and then incubated with 250ng ARG-seq or eRNA-seq RNA baits overnight at 65C in hybridization buffer (2.5ug Cot1 DNA (ThermoFisher), 2.5ug Salmon Sperm DNA (ThermoFisher), 15mM p5 blocking primers, 15mM p7 blocking primers, 5X SSPE (ThermoFisher), 5X Denhardt’s Solution (ThermoFisher), 0.133% SDS). Blocking primers are: p5-AATGATACGGCGACCACCGAGATCTACAC, ACACTCTTTCCCTACACGACGCTCTTCCGATC/3InvdT/ p7-CAAGCAGAAGACGGCATACGAGAT, GTGACTGGAGTTCAGACGTGT GCTCTTCCGATC/3InvdT/ Primers for amplification are: p5-AATGATACGGCGACCACCGAGA, p7-CAAGCAGAAGACGGCATACGAG.

Hybridized samples were incubated with MyOne Streptavadin T1 Dynabeads (Invitrogen) in binding buffer (1M NaCl, 10mM Tris-HCl pH 7.5, 1mM EDTA). Beads were washed once in 1x SCC, 0.1% SDS at room temperature and three times in 0.1x SCC 0.1% SDS at 65C. Captured libraries were eluted with 0.1M NaOH and neutralized with 1M Tris-HCl pH 7.5. Libraries were then purified using the Qiagen MinElute PCR cleanup kit and re-amplified using Herculase II Fusion polymerase (Agilent). Sequencing for mRNAseq was at a depth of 3-5 million reads per library and for eRNAseq, 20-30 million reads per library on a NextSeq.

Reads were aligned and processed as with total RNA-seq, and raw count data was analyzed using R, including edgeR (Robinson et al., 2009). Data was also normalized by the geometric mean of the reads from control genes or enhancers. Control regions were identified as regions that do not change with KCl in published RNA-seq data (Kim et al., 2010).

### Western blotting

To detect protein expression in mouse cortical neurons, neurons were collected in cold lysis buffer: 1% Triton X-100, 50 mM HEPES, pH 7.4, 150 mM NaCl, 1.5 mM MgCl_2_, 1 mM EGTA, 10% glycerol, and freshly added protease and phosphatase inhibitors from Roche Applied Science Cat. # 05056489001 and 04906837001, respectively. Lysed neurons were treated with 4X sample buffer (40% glycerol, 8% SDS, 0.25M Tris-HCL, pH 6.8, 10% 2-mercaptoethanol) and boiled for 5 minutes. Samples were centrifuged at full speed for 3 minutes before loading on NuPage 4-12% Bis-Tris Gels (Invitrogen). Gels were run at 140V for 55 minutes. We transferred onto nitrocellulose membranes using the BioRad transfer system at 114V for 1h and 7min. Membranes were blocked in 5% milk-TBST for 1 hour. They were treated with primary antibody in 5% milk-TBST for at least one hour at room temperature or overnight at 4C. To visualize protein, blots were incubated with secondary antibody in TBST in the dark for 45 minutes. Blots were imaged using a LiCor Odyessy and quantified using ImageJ. Primary antibodies used were: rabbit anti-phosphoERK1/2 (Cell Signaling Technology 4370, 1:1000), mouse anti-GAPDH (Pierce, GA1R, 1:10000), rabbit anti-ARC (Synaptic Systems, 156-003, 1:1000). Secondary antibodies used were: IDR dye 680 goat anti-rabbit (LiCor, 1:10000), IDR dye 800 goat anti-mouse (LiCor, 1:10000).

To detect protein expression in rat cortical neurons, neurons were disrupted by brief sonication (three cycles of 30 sec in low setting in Bioruptor at 4 degrees C) and then cleared of debris by high-speed centrifugation (14500 RPM for 1 minute). The supernatant was collected in separate tubes and resolved by gel electrophoresis on 4-20% pre-cast gels (Life technology) and transferred to a nitrocellulose membrane using the iBlot gel transfer apparatus (Life technology). Immunoblots were incubated with primary antibody overnight. Blots were visualized with a LiCor Odyssey infrared scanner after immunolabeling primary antibodies with infrared fluorophore-tagged secondary antibody (Molecular Probes). Images were processed using the Odyssey 2.1 software. Primary antibodies used were: rabbit anti-phosphoERK1/2 (Cell Signaling Technology 4370), H4 (Cell Signaling Technology 2935), Actin (Millipore, AM4302).

### Nuclear isolation for western blotting

Nuclear lysate was prepared from treated neurons by first liberating the nuclei in a non-ionic detergent buffer [10mM HEPES (pH 7.9), 10mM KCl, 2mM MgCl_2_, 0.5mM dithiothretol, 0.1% NP-40] for precisely 30 seconds and subsequently lysing them in NETN buffer [0.5% NP-40, 1mM EDTA, 50mM Tris, 120mM NaCl, pH 7.5] freshly supplemented with 0.5% protease inhibitor cocktail (Sigma) and phosphatase inhibitor cocktails (Sigma). Nuclear liberation was confirmed under the microscope before the released nuclei was scraped and dissolved in the NETN buffer (Saha et al., 2009).

### Chromatin immunoprecipitation (ChIP)

Media on the neurons was removed and neurons were fixed in crosslinking buffer (10 mM HEPES pH 7.5, 100 mM NaCl, 1 mM EDTA, 1 mM EGTA, 1% formaldehyde) for ten minutes at room temperature, and this reaction was quenched using 125mM glycine for 5 minutes. For H3K27ac ChIP, 250,000 neurons were used per ChIP sample. For Pol2 ChIP, 2 million neurons were used per sample. Neurons were then washed with cold PBS and then collected in PBS with 0.25% BSA and pelleted by centrifuging at 700 x g for 15 minutes. Cell pellets were stored at ˗80C. Neurons were sonicated using a Covaris E3 sonicator in lysis buffer (10 mM Tris pH 8.0, 1mM EDTA, 1 mM EGTA, 1X Roche complete EDTA-free protease inhibitors, 0.15% SDS). Sonication was done for 8 minutes per samples with 200 cycles/burst, a 2% duty cycle at power level 3. This reliably produced fragments between 100 and 700bp in length. Samples were then supplemented with ChIP Buffer to make SDS-ChIP buffer (10 mM Tris pH 8.0, 0.1% SDS,1% Triton X-100, 150 mM NaCl, 1 mM EDTA, 0.3 mM EGTA, 1X Roche complete EDTA-free protease inhibitors). For H3K27ac ChIP, Protein A beads (Dynabeads) were washed with 1% BSA/TBST and added to the fragmented DNA for a pre-clear and rotated at 4C for one hour. A different set of protein A beads was pre-treated with .48ug of antibody (Abcam, ab4729)/experiment for H3K27ac ChIP. The same procedure was followed for Pol2 ChIP, but with Protein G Dynabeads and 4ug antibody (Abcam, ab817) per crosslinked input. Following the pre-clear, pre-clear beads were removed, an aliquot of fragmented DNA was set aside as the input, and antibody-treated beads were incubated with the fragmented DNA overnight at 4C. were washed twice with cold low salt wash buffer (0.1% SDS, 20 mM Tris pH 8.0, 1% Triton X-100, 150 mM NaCl, 2 mM EDTA), twice with cold high salt wash buffer (0.1% SDS, 20 mM Tris pH 8.0, 1% Triton X-100, 500 mM NaCl, 2 mM EDTA), twice with cold LiCl wash buffer (1% NaDOC, 10 mM Tris pH 8.0, 1% NP40, 250 mM LiCl, 1 mM EDTA), and once with room temperature TE. Crosslinks were reversed by incubating samples in TE+1%SDS at 65C overnight. Samples were then treated with RNAse A and Proteinase K, and DNA was eluted using MinElute Columns (Qiagen) according to the instructions of the manufacturer.

### Chromatin immunoprecipitation sequencing (ChIP-seq)

For H3K27ac ChIP-seq, libraries were prepared using 5ug of immunoprecipitated DNA or input DNA with the NuGen Ultralow V2 1-96 library prep kit. Libraries were sequenced on an Illumina NextSeq to a depth of at least 30 million reads per library. Reads were aligned to mouse genome mm9 using bowtie2 (Langmead and Salzberg, 2012). The resulting SAM files were made into BED files using SAMtools and BEDtools, with reads extended to 300 base pairs (Li et al., 2009; Quinlan and Hall, 2010) and then into bigWig files using UCSC-tools (Kuhn et al., 2013). Reads were assigned to individual enhancers or promoters using bedtools map and data was analyzed using R.

For downstream analysis, H3K27ac ChIP-seq data was input-normalized and then normalized by dividing by the geometric mean of control enhancers identified based on their location near the same control genes used for ARG-seq (control enhancer selection described in Capture RNA sequencing section). We confirmed using the Tukey HSD test in conjunction with ANOVA that read-depth-normalized signal at control enhancers was not affected by stimulation or by addition of U0126. We also performed one replicate using *Drosophila* spike-in chromatin (Active Motif #61686, #53083) according to the instructions of the manufacturer and observed that U0126 treatment did not result in global H3K27ac changes. The data plotted in the figures shows the mean input-normalized and control-normalized signal from the same regions targeted by eRNA-seq of each enhancer for two biological replicates, averaging each enhancer across replicates prior to plotting, and including only enhancers captured in eRNA-seq.

For Pol2 ChIP-seq, reads were aligned to mouse genome mm9 using the STAR aligner (Dobin et al., 2013). The resulting SAM files were made into read-extended (200 bases per fragment) BED files using SAMtools and BEDtools (Li et al., 2009; Quinlan and Hall, 2010) and then into bigWig files using UCSC-tools (Kuhn et al., 2013). For analysis, the metaseq (Dale et al., 2014), numpy (Van Der Walt et al., 2011), and matplotlib (Hunter, 2007) python packages were used to process aligned bam files, extend reads to 200 bases, and to produce read-depth- and input-normalized data. Additional analysis was performed in R. Given across-sample variability in read-depth- and input-normalized data, the samples were further normalized to RNAPII ChIP-seq density measured at constitutively active, non-activity-regulated control gene promoters—similar to the across-sample ChIP-seq normalization methods adopted by others for quantitative analysis of peaks (Shao et al., 2012). Specifically, data from each sample was normalized to the median value of a distribution of RNAPII density values occurring at ~800 constitutively active TSSs (˗300 to +300bp) with unchanging mRNA levels under KCl as measured by RNAseq (Kim, et al. 2011).

For analysis of published data, data from Kim et al. 2010 was used as aligned and processed by the authors. Data from Telese et al. 2014 was downloaded from GEO as fastq files, re-aligned to mm9, and processed as above. Data from ENCODE was downloaded as processed by authors. Signal was binned across TSSs and enhancers using the Python package metaseq (R. K. Dale et al., 2014). Plots were made using R, smoothing with the lowess function.

### Luciferase assays

The sequences for enhancer e5 was amplified using PCR from genomic DNA extracted from wildtype (C57BL/6J) mice, utilizing primers that included flanking KpnI and XhoI sites (ATACGGTACCCGAGACTACGTCA, ATGTCTCGAGATTAAAAAGGCCC). These amplified sequences were cloned into pTAN02, an ITR-containing AAV screening vector containing minimal human pFos upstream of the Firefly luciferase gene (Nguyen at al., 2016) with the KpnI and XhoI sites. Additionally, pTAN02 without an enhancer insert was included as a “no enhancer” control. Primary cortical neuron cultures (see above) were transfected using PEI (4:1 PEI:DNA mass ratio) on DIV5. These cultures were co-transfected with an internal control Renilla luciferase construct, pTK-RN, at a fixed mass ratio of 9:1, Firefly construct:Renilla construct. Each experiment was run in triplicate. 30 minutes prior to depolarization, 10uM U0126 in DMSO or a DMSO vehicle was added to the culture media. Cultures were depolarized by changing media to complete neurobasal with 55mM KCl and were incubated for 12 hours. A non-depolarized control received a media change with no additional KCl. Cultures were collected on the night of DIV7 and prepared using the Dual-Luciferase Reporter Assay System (Promega) according to the manufacturer’s protocol. The lysate was assayed over a 10 second period using the GloMax 20/20 Single Tube Luminometer (Promega), and the luciferase activity was calculated as a ratio of the Firefly to Renilla output values.

## QUANTIFICATION AND STATISTICAL ANALYSIS

We have included most statistical details in our Figure legends, including p-values, statistical tests used, ‘n’s for each experiment, and a description of to what ‘n’ refers. Biological replicates refer to biological material from different mice (all experiments), with biological replicate samples also collected on a different day (in vitro experiments only).

### Classification of rapid PRGs, delayed PRGs and SRGs

In experiments in mouse cortical neurons, We classified genes as rapid PRGs, delayed PRGs, or SRGs based on a comparison of expression at 6h of sustained KCl depolarization with and without cycloheximide treatment, as well as the kinetics of induction (main text). We eliminated four PRGs from our analysis due to ambiguity in our classification scheme, which exclusively relied upon kinetics of induction to distinguish rapid from delayed PRGs. We eliminated two genes (*Vgf* and *Homer1*) because their expression peaked at 6 hours of KCl stimulus, but they showed robust and significant pre-mRNA induction at 20 minutes. We also eliminated two genes (*Gadd45b* and *Nfkbid*) because while their mRNA induction peaked at 1h, they did not show a trend towards pre-mRNA or mRNA induction at 20 minutes of KCl. For significance testing in the classification, we used edgeR’s glmFit and glmTreat functions (Robinson et al., 2009). PCA was performed using the prcomp function in R using normalized mRNA expression values. Specifically, to better assess expression kinetics, each gene was normalized such that its lowest expression value was set at 1 and its highest at 10.

For in vivo data, gene classification was based on in vitro mouse data. However, we eliminated delayed PRGs with higher induction at 30 minutes compared to 150 minutes of visual stimulus.

### Nearest-neighbor classifier

Our classifier for post-hoc determination of activity pattern based on gene expression used the maximum expression at any time point for each gene, such that the kinetics of gene induction did not contribute to the classifier. It compared each replicate in a testing set to all replicates in a training set using Euclidean distance, and classified based on the minimum distance. It was run with both separate testing and training sets (6 biological replicates each) and leave-one-out cross validation.

### RNA-seq

For ARG-seq and total RNA-seq figures, we plotted a mean of the control-normalized expression levels for each gene from several biological replicates. All p-values reported in the figure legends for comparisons between two groups of genes are from a non-parametric two-tailed Wilcoxon rank-sum test (unless otherwise noted). A paired test was used when comparing between the same set of genes in two conditions. We confirmed significance using a two-tailed Student’s T-test. We also confirmed that the differences observed via analysis of the mean expression levels were replicated in each biological replicate individually (p<0.05, rank-sum test).

For ARG-seq and eRNA-seq, we confirmed using the Tukey HSD test in conjunction with ANOVA that expression from control genes or control enhancers in read-depth-normalized samples and spike-in-normalized samples is not affected by membrane depolarization, visual stimulation, or addition of U0126/SL327.

### Enhancer transcription factor and modified histone binding

For the enhancer data, in addition to the data shown in the figures, we also compared only those rapid and delayed enhancers near delayed PRGs. In unstimulated neurons, for SRF, CREB, MEF2, MED23, MED1 and NCoR we compared binding ˗6kb to +6kb from the centers of rapid enhancers compared to delayed enhancers and as reported in the main text found greater binding at rapid enhancers (p<0.009, rank-sum test, including only enhancers within 100 kb of delayed PRGs). Active histone marks H3K27ac, H3K4me2, H3K4me1, and H4K16ac were also higher in a comparison of the same rapid compared to delayed enhancers in unstimulated neurons (p<0.01, rank-sum test, only enhancers within 100 kb of delayed PRGs).

### H3K27ac ChIP-seq

For the H3K27ac ChIP-seq, all p-values reported are from the two-tailed non-parametric Wilcoxon rank-sum test, but we confirmed significance using the Student’s t-test. We also performed a Student’s *t*-test comparing the mean signal across all enhancers from each replicate for each gene class without U0126 to the mean signal across enhancers from each gene class with U0126 and found no significant difference (p>0.6). We also compared each enhancer individually, and again found no significant change in H3K27ac signal at any enhancer with U0126 (p>0.9, Bonferroni corrected).

### Pol2 ChIP-seq

Additional analysis was performed in R. Given across-sample variability in read-depth- and input-normalized data, the samples were further normalized to RNAPII ChIP-seq density measured at constitutively active, non-activity-regulated control gene promoters—similar to the across-sample ChIP-seq normalization methods adopted by others for quantitative analysis of peaks (Shao et al., 2012). Specifically, data from each sample was normalized to the median value of a distribution of RNAPII density values occurring at ~800 constitutively active TSSs (˗300 to +300bp) with unchanging mRNA levels under KCl as measured by RNAseq (Kim, et al. 2011). As a separate analysis, early and delayed TSS lists were filtered for TSS’s with mean Pol2 ChIPseq density greater than a threshold condition defined as two standard deviations above the mean value of un-expressed (Kim, et al. 2011) negative control TSS’.

## DATA AND SOFTWARE AVAILABILITY

The RNA-Seq and ChIP-Seq data generated in this work are in-process for GEO submission. In the meantime, processed data for RNA-Seq is available Table S2.

## Supplemental table legends

**Table S1. Targeted capture oligonucleotides.** Oligonucleotides used to make biotinylated probes for ARG-seq and eRNA-seq. Sequences include in-vitro-transcription promoters and amplification primer sites.

Related to Figure1, Figure S1A, Figure 7

**Table S2. ARG-seq and eRNA-seq data.** Data from each replicate of each experiment normalized for read depth and by controls, as in methods.

Related to Figure 1, S1, Figure 3, S3, Figure 4, Figure 6, S6, Figure 7

## REFERENCES

Abbott, L.F., Nelson, S.B., 2000. Synaptic plasticity: taming the beast. Nat. Neurosci. 3, 1178–1183.

Adelman, K., Lis, J.T., 2012. Promoter-proximal pausing of RNA polymerase II: emerging roles in metazoans. Nat. Rev. Genet. 13, 720–731. doi: http://dx.doi.org/10.7554/eLife.02407

Allen, B.L., Taatjes, D.J., 2015. The Mediator complex: a central integrator of transcription. Nat. Rev. Mol. Cell Biol. 16, 155–166. doi:10.1038/nrm3951

Arnold, F.J.L., Hofmann, F., Bengtson, C.P., Wittmann, M., Vanhoutte, P., Bading, H., 2005. Microelectrode array recordings of cultured hippocampal networks reveal a simple model for transcription and protein synthesis-dependent plasticity. J. Physiol. 564, 3–19. doi:10.1113/jphysiol.2004.077446

Atkins, C.M., Selcher, J.C., Petraitis, J.J., Trzaskos, J.M., Sweatt, J.D., 1998. The MAPK cascade is required for mammalian associative learning. Nat. Neurosci. 1.

Besnard, A., Galan-Rodriguez, B., Vanhoutte, P., Caboche, J., 2011. Elk-1 a transcription factor with multiple facets in the brain. Front. Neurosci. 5, 1–11. doi:10.3389/fnins.2011.00035

Bollen, M., Peti, W., Ragusa, M.J., Beullens, M., 2010. The extended PP1 toolkit: Designed to create specificity. Trends Biochem. Sci. doi:10.1016/j.tibs.2010.03.002

Bose, D.A., Donahue, G., Reinberg, D., Shiekhattar, R., Bonasio, R., Berger, S.L., 2017. RNA Binding to CBP Stimulates Histone Acetylation and Transcription. Cell 168, 135–149.e22. doi:10.1016/j.cell.2016.12.020

Chepelev, I., Wei, G., Wangsa, D., Tang, Q., Zhao, K., 2012. Characterization of genome-wide enhancer-promoter interactions reveals co-expression of interacting genes and modes of higher order chromatin organization. Cell Res. 22, 490–503. doi:10.1038/cr.2012.15

Cho, J.-H., Huang, B.S., Gray, J.M., 2016. RNA sequencing from neural ensembles activated during fear conditioning in the mouse temporal association cortex. Sci. Rep. 6, 31753. doi:10.1038/srep31753

Chowdhury, S., Shepherd, J.D., Okuno, H., Lyford, G., Petralia, R.S., Plath, N., Kuhl, D., Huganir, R.L., Worley, P.F., 2006. Arc/Arg3.1 Interacts with the Endocytic Machinery to Regulate AMPA Receptor Trafficking. Neuron 52, 445–459. doi:10.1016/j.neuron.2006.08.033

Churchman, L.S., Weissman, J.S., 2011. Nascent transcript sequencing visualizes transcription at nucleotide resolution. Nature 469, 368–73. doi:10.1038/nature09652

ENCODE Project Consortium et al., 2012. An integrated encyclopedia of DNA elements in the human genome. Nature 489, 57–74. doi:10.1038/nature11247

Core, L.J., Waterfall, J.J., Lis, J.T., 2008. Nascent RNA sequencing reveals widespread pausing and divergent initiation at human promoters. Science 322, 1845–8. doi:10.1126/science.1162228

Creyghton, M.P., Cheng, A.W., Welstead, G.G., Kooistra, T., Carey, B.W., Steine, E.J., Hanna, J., Lodato, M. a, Frampton, G.M., Sharp, P. a, Boyer, L. a, Young, R. a, Jaenisch, R., 2010. Histone H3K27ac separates active from poised enhancers and predicts developmental state. Proc. Natl. Acad. Sci. U. S. A. 107, 21931–21936. doi:10.1073/pnas.1016071107

Davis, S., Vanhoutte, P., Pages, C., Caboche, J., Laroche, S., 2000. The MAPK / ERK Cascade Targets Both Elk-1 and cAMP Response Element-Binding Protein to Control Long-Term Potentiation-Dependent Gene Expression in the Dentate Gyrus In Vivo. J. Neurosci. 20,4563–4572.

De Koninck, P., Schulman, H., 1998. Sensitivity of CaM Kinase II to the Frequency of Ca2+ Oscillations. Science (80-.). 279, 227–230. doi:10.1126/science.279.5348.227

Dolmetsch, R., Xu, K., Lewis, R.S., 1998. Calcium oscillations increase the efficiency and specificity of gene expression. Nature 392, 933–936.

Dolmetsch, R.E., Lewis, R.S., Goodnow, C.C., Healy, J.I., 1997. Differential activation of transcription factors induced by calcium response amplitude and duration. Nature.

Dolmetsch, R.E., Pajvani, U., Fife, K., Spotts, J.M., Greenberg, M.E., 2001. Signaling to the nucleus by an L-type calcium channel-calmodulin complex through the MAP kinase pathway. Science 294, 333–9. doi:10.1126/science.1063395

Douglas, R.M., Dragunow, M., Robertson, H.A., 1988. High-frequency discharge of dentate granule cells, but not long-term potentiation, induces c-fos protein. Mol. Brain Res. 4, 259–262.

Dudek, S.M., Fields, R.D., 2001. Mitogen-Activated Protein Kinase / Extracellular Signal-Regulated Kinase Activation in Somatodendritic Compartments: Roles of Action Potentials, Frequency, and Mode of Calcium Entry. J. Neurosci. 21, 1–5.

El-Sherif, E., Levine, M., 2016. Shadow Enhancers Mediate Dynamic Shifts of Gap Gene Expression in the Drosophila Embryo. Curr. Biol. 26, 1164–1169. doi:10.1016/j.cub.2016.02.054

Eriksson, M., Taskinen, M., Leppä, S., 2006. Mitogen Activated Protein Kinase-Dependent Activation of c-Jun and c-Fos is required for Neuronal differentiation but not for Growth and Stress Reposne in PC12 cells. J. Cell. Physiol. 207, 12–22. doi:10.1002/JCP

Escoubet-Lozach, L., Benner, C., Kaikkonen, M.U., Lozach, J., Heinz, S., Spann, N.J., Crotti, A., Stender, J., Ghisletti, S., Reichart, D., Cheng, C.S., Luna, R., Ludka, C., Sasik, R., Garcia-Bassets, I., Hoffmann, A., Subramaniam, S., Hardiman, G., Rosenfeld, M.G., Glass, C.K., 2011. Mechanisms establishing tlr4-responsive activation states of inflammatory response genes. PLoS Genet. 7. doi:10.1371/journal.pgen.1002401

Eshete, F., Fields, R.D., 2001. Spike Frequency Decoding and Autonomous Activation of Ca 2+–Calmodulin-Dependent Protein Kinase II in Dorsal Root Ganglion Neurons. J. Neurosci. 21,6694–6705.

Esnault, C., Gualdrini, F., Horswell, S., Kelly, G., Stewart, A., East, P., Matthews, N., Treisman, R., 2017. ERK-Induced Activation of TCF Family of SRF Cofactors Initiates a Chromatin Modification Cascade Associated with Transcription. Mol. Cell 65, 1081–1095.e5. doi:10.1016/j.molcel.2017.02.005

Evans, M.D., Sammons, R.P., Lebron, S., Dumitrescu, A.S., Watkins, T.B.K., Uebele, V.N., Renger, J.J., Grubb, M.S., 2013. Calcineurin signaling mediates activity-dependent relocation of the axon initial segment. J. Neurosci. 33, 6950–63. doi:10.1523/JNEUROSCI.0277-13.2013

Favata, M.F., Horiuchi, K.Y., Manos, E.J., Daulerio, A.J., Stradley, D.A., Feeser, W.S., Van Dyk, D.E., Pitts, W.J., Earl, R.A., Hobbs, F., Copeland, R.A., Magolda, R.L., Scherle, P.A., Trzaskos, J.M., 1998. Identification of a novel inhibitor of mitogen-activated protein kinase kinase. J. Biol. Chem. 273, 18623–32. doi:10.1074/JBC.273.29.18623

Fenouil, R., Cauchy, P., Koch, F., Descostes, N., Cabeza, J.Z., Innocenti, C., Ferrier, P., Spicuglia, S., Gut, M., Gut, I., Andrau, J.C., 2012. CpG islands and GC content dictate nucleosome depletion in a transcription-independent manner at mammalian promoters. Genome Res. 22, 2399–2408. doi:10.1101/gr.138776.112

Fields, R.D., Eshete, F., Stevens, B., Itoh, K., 1997. Action Potential-Dependent Regulation of Gene Expression: Temporal Specificity in Ca 2+, cAMP-Responsive Element Binding Proteins, and Mitogen-Activated Protein Kinase Signaling. J. Neurosci. 17, 7252–7266.

Fields, R.D., Lee, P.R., Cohen, J.E., 2005. Temporal integration of intracellular Ca2+ signaling networks in regulating gene expression by action potentials. Cell Calcium 37, 433–42. doi:10.1016/j.ceca.2005.01.011

Flavell, S.W., Greenberg, M.E., 2008. Signaling mechanisms linking neuronal activity to gene expression and plasticity of the nervous system. Annu. Rev. Neurosci. 31, 563–90. doi:10.1146/annurev.neuro.31.060407.125631

Fowler, T., Sen, R., Roy, A.L., 2011. Regulation of primary response genes. Mol. Cell 44, 348–360. doi:10.1016/j.molcel.2011.09.014

Frank, C.L., Liu, F., Wijayatunge, R., Song, L., Biegler, M.T., Yang, M.G., Vockley, C.M., Safi, A., Gersbach, C.A., Crawford, G.E., West, A.E., 2015. Regulation of chromatin accessibility and Zic binding at enhancers in the developing cerebellum. Nat. Neurosci. 18, 647–656. doi:10.1038/nn.3995

Fujii, H., Inoue, M., Okuno, H., Sano, Y., Takemoto-Kimura, S., Kitamura, K., Kano, M., Bito, H., 2013. Nonlinear decoding and asymmetric representation of neuronal input information by CaMKIIa and calcineurin. Cell Rep. 3, 978–87. doi:10.1016/j.celrep.2013.03.033

Gaidatzis, D., Burger, L., Florescu, M., Stadler, M.B., 2015. Analysis of intronic and exonic reads in RNA-seq data characterizes transcriptional and post-transcriptional regulation. Nat Biotech 33, 722–729. doi:10.1038/nbt.3269

Gilchrist, D.A., Dos Santos G., Fargo, D.C., Xie, B., Gao, Y., Li, L., Adelman, K., 2010. Pausing of RNA polymerase II disrupts DNA-specified nucleosome organization to enable precise gene regulation. Cell 143, 540–551. doi:10.1016/j.cell.2010.10.004

Gotoh, Y., Nishida, E., Yamashita, T., Hoshi, M., Kawakami, M., Sakai, H., 1990. Microtubule-associated-protein (MAP) kinase activated by nerve growth factor and epidermal growth factor in PC12 cells. Identity with the mitogen-activated MAP kinase of fibroblastic cells.Eur. J. Biochem. 193, 661–9.

Gray, J.M., Harmin, D.A., Boswell, S.A., Cloonan, N., Mullen, T.E., Ling, J.J., Miller, N., Kuersten, S., Ma, Y.C., McCarroll, S.A., Grimmond, S.M., Springer, M., 2014. SnapShot-Seq: A method for extracting genome-wide, in Vivo mRNA dynamics from a single total RNA sample. PLoS One 9. doi:10.1371/journal.pone.0089673

Greenberg, M.E., Ziff, E.B., Greene, L.A., 1986. Stimulation of Neuronal Acetylcholine Receptors Induces Rapid Gene Transcription. Science 234, 80–83.

Gualdrini, F., Esnault, C., Horswell, S., Stewart, A., Matthews, N., Treisman, R., 2016. SRF Cofactors Control the Balance between Cell Proliferation and Contractility. Mol. Cell 1–14. doi:10.1016/j.molcel.2016.10.016

Guan, J., Haggarty, S.J., Giacometti, E., Dannenberg, J., Joseph, N., Gao, J., Nieland, T.J.F., Zhou, Y., Wang, X., Mazitschek, R., Bradner, J.E., Depinho, R.A., Jaenisch, R., Tsai, L., 2009. HDAC2 negatively regulates memory formation and synaptic plasticity. Nature 459, 55–60. doi:10.1038/nature07925

Hah, N., Murakami, S., Nagari, A., Danko, C.G., Kraus, W.L., 2013. Enhancer transcripts mark active estrogen receptor binding sites Enhancer transcripts mark active estrogen receptor binding sites. Genome Res. 23, 1210–1223. doi:10.1101/gr.152306.112

Hardingham, G.E., Arnold, F.J.L., Bading, H., 2001. A calcium microdomain near NMDA receptors: on switch for ERK-dependent communication. Nat. Neurosci. 12–13.

Hargreaves, D.C., Horng, T., Medzhitov, R., 2009. Control of Inducible Gene Expression by Signal-Dependent Transcriptional Elongation. Cell 138, 129–145. doi:10.1016/j.cell.2009.05.047

Herschman, H.R., 1991. Primary Response Genes Induced by Growth Factors and Tumor Promoters. Annu. Rev. Biochem. 60, 281–319.

Huang, Y.Y., Martin, K.C., Kandel, E.R., 2000. Both protein kinase A and mitogen-activated protein kinase are required in the amygdala for the macromolecular synthesis-dependent late phase of long-term potentiation. J. Neurosci. 20, 6317–6325. doi:20/17/6317 [pii]

Ibata, K., Sun, Q., Turrigiano, G.G., 2008. Rapid synaptic scaling induced by changes in postsynaptic firing. Neuron 57, 819–26. doi:10.1016/j.neuron.2008.02.031

Jindal, G.A., Goyal, Y., Burdine, R.D., Rauen, K.A., Shvartsman, S.Y., 2015. RASopathies: Unraveling mechanisms with animal models. DMM Dis. Model. Mech. 8, 769–782. doi:10.1242/dmm.020339

Joo, J.-Y., Schaukowitch, K., Farbiak, L., Kilaru, G., Kim, T.-K., 2015. Stimulus-specific combinatorial functionality of neuronal c-fos enhancers. Nat. Neurosci. 1–12. doi:10.1038/nn.4170

Josefowicz, S.Z., Shimada, M., Armache, A., Li, C.H., Miller, R.M., Lin, S., Yang, A., Dill, B.D., Molina, H., Park, H.S., Garcia, B.A., Taunton, J., Roeder, R.G., Allis, C.D., 2016. Chromatin Kinases Act on Transcription Factors and Histone Tails in Regulation of Inducible Transcription. Mol. Cell 64, 347–361. doi:10.1016/j.molcel.2016.09.026

Kaikkonen, M.U., Spann, N., Heinz, S., Romanoski, C.E., Karmel, A., Stender, J.D., Chun, H.B., Tough, D.F., Prinjha, R.K., Benner, C., Glass, C.K., 2013. Activation Is Coupled To Enhancer Transcription. Mol. Cell 51, 310–325. doi:10.1016/j.molcel.2013.07.010.Remodeling

Kholodenko, B.N., Hancock, J.F., Kolch, W., 2010. Signalling ballet in space and time. Nat. Rev. Mol. Cell Biol. 11, 414–26. doi:10.1038/nrm2901

Kim, T.-K., Hemberg, M., Gray, J.M., Costa, A.M., Bear, D.M., Wu, J., Harmin, D. a, Laptewicz, M., Barbara-Haley, K., Kuersten, S., Markenscoff-Papadimitriou, E., Kuhl, D., Bito, H., Worley, P.F., Kreiman, G., Greenberg, M.E., 2010. Widespread transcription at neuronal activity-regulated enhancers. Nature 465, 182–187. doi:10.1038/nature09033

Kingsbury, T.J., Bambrick, L.L., Roby, C.D., Krueger, B.K., 2007. Calcineurin activity is required for depolarization-induced, CREB-dependent gene transcription in cortical neurons. J. Neurochem. 103, 761–770. doi:10.1111/j.1471-4159.2007.04801.x

Korb, E., Herre, M., Zucker-Scharff, I., Darnell, R.B., Allis, C.D., 2015. BET protein Brd4 activates transcription in neurons and BET inhibitor Jq1 blocks memory in mice. Nat. Neurosci. 1–13. doi:10.1038/nn.4095

Kuja-Panula, J., Kiiltomäki, M., Yamashiro, T., Rouhiainen, A., Rauvala, H., 2003. AMIGO, a transmembrane protein implicated in axon tract development, defines a novel protein family with leucine-rich repeats. J. Cell Biol. 160, 963–973. doi:10.1083/jcb.200209074

Li, X., Zhou, W., Zeng, S., Liu, M., Luo, Q., 2007. Long-term recording on multi-electrode array reveals degraded inhibitory connection in neuronal network development. Biosens. Bioelectron. 22, 1538–1543. doi:10.1016/j.bios.2006.05.030

Ma, H., Groth, R.D., Wheeler, D.G., Barrett, C.F., Tsien, R.W., 2011. Excitation-transcription coupling in sympathetic neurons and the molecular mechanism of its initiation. Neurosci. Res. 70, 2–8. doi:10.1016/j.neures.2011.02.004

Marais, R., Wynne, J., Treisman, R., 1993. The SRF accessory protein Elk-1 contains a growth factor-regulated transcriptional activation domain. Cell 73, 381–393. doi:10.1016/0092-8674(93)90237-K

Mardinly, A.R., Spiegel, I., Patrizi, A., Centofante, E., Bazinet, J.E., Tzeng, C.P., Mandel-Brehm, C., Harmin, D.A., Adesnik, H., Fagiolini, M., Greenberg, M.E., 2016. Sensory experience regulates cortical inhibition by inducing IGF1 in VIP neurons. Nature 531, 371–375. doi:10.1038/nature17187

Marshall, C.J., 1995. Specificity of receptor tyrosine kinase signaling: transient versus sustained extracellular signal-regulated kinase activation. Cell 80, 179–185. doi:10.1016/0092-8674(95)90401-8

Maze, I., Wenderski, W., Noh, K.M., Bagot, R.C., Tzavaras, N., Purushothaman, I., Elsässer, S.J., Guo, Y., Ionete, C., Hurd, Y.L., Tamminga, C.A., Halene, T., Farrelly, L., Soshnev, A.A., Wen, D., Rafii, S., Birtwistle, M.R., Akbarian, S., Buchholz, B.A., Blitzer, R.D., Nestler, E.J., Yuan, Z.F., Garcia, B.A., Shen, L., Molina, H., Allis, C.D., 2015. Critical Role of Histone Turnover in Neuronal Transcription and Plasticity. Neuron 87, 77–94. doi:10.1016/j.neuron.2015.06.014

Mercer, T.R., Gerhardt, D.J., Dinger, M.E., Crawford, J., Trapnell, C., Jeddeloh, J. a, Mattick, J.S., Rinn, J.L., 2011. Targeted RNA sequencing reveals the deep complexity of the human transcriptome. Nat. Biotechnol. 30, 99–104. doi:10.1038/nbt.2024

Murphy, L.O., MacKeigan, J.P., Blenis, J., 2004. A Network of Immediate Early Gene Products Propagates Subtle Differences in Mitogen-Activated Protein Kinase Signal Amplitude and Duration. Mol. Cell. Biol. 24, 144–153. doi:10.1128/MCB.24.1.144-153.2004

Murphy, L.O., Smith, S., Chen, R.-H., Fingar, D.C., Blenis, J., 2002. Molecular interpretation of ERK signal duration by immediate early gene products. Nat. Cell Biol. 4, 556–64. doi:10.1038/ncb822

Murphy, T.H., Blatter, L. a, Bhat, R. V, Fiore, R.S., Wier, W.G., Baraban, J.M., 1994. Differential regulation of calcium/calmodulin-dependent protein kinase II and p42 MAP kinase activity by synaptic transmission. J. Neurosci. 14, 1320–1331.

Nguyen, P. V, Abel, T., Kandel, E.R., 1994. Requirement of a critical period of transcription for induction of a late phase of LTP. Science 265, 1104–7.

Nicodeme, E., Jeffrey, K.L., Schaefer, U., Beinke, S., Dewell, S., Chung, C., Chandwani, R., Marazzi, I., Wilson, P., Coste, H., White, J., Kirilovsky, J., Rice, C.M., Lora, J.M., Prinjha, R.K., Lee, K., Tarakhovsky, A., 2010. Suppression of inflammation by a synthetic histone mimic. Nature 468, 1119–1123. doi:10.1038/nature09589

Nord, A.S., Blow, M.J., Attanasio, C., Akiyama, J.A., Holt, A., Hosseini, R., Phouanenavong, S., Plajzer-Frick, I., Shoukry, M., Afzal, V., Rubenstein, J.L.R., Rubin, E.M., Pennacchio, L.A., Visel, A., 2013. Rapid and pervasive changes in genome-wide enhancer usage during mammalian development. Cell 155, 1521–1531. doi:10.1016/j.cell.2013.11.033

Ostuni, R., Piccolo, V., Barozzi, I., Polletti, S., Termanini, A., Bonifacio, S., Curina, A., Prosperini, E., Ghisletti, S., Natoli, G., 2013. Latent enhancers activated by stimulation in differentiated cells. Cell 152, 157–171. doi:10.1016/j.cell.2012.12.018

Plath, N., Ohana, O., Dammermann, B., Errington, M.L., Schmitz, D., Gross, C., Mao, X., Engelsberg, A., Mahlke, C., Welzl, H., Kobalz, U., Stawrakakis, A., Fernandez, E., Waltereit, R., Bick-Sander, A., Therstappen, E., Cooke, S.F., Blanquet, V., Wurst, W., Salmen, B., Bösl, M.R., Lipp, H.P., Grant, S.G.N., Bliss, T.V.P., Wolfer, D.P., Kuhl, D., 2006. Arc/Arg3.1 Is Essential for the Consolidation of Synaptic Plasticity and Memories. Neuron 52, 437–444. doi:10.1016/j.neuron.2006.08.024

Rada-Iglesias, A., Bajpai, R., Swigut, T., Brugmann, S. a, Flynn, R. a, Wysocka, J., 2011. A unique chromatin signature uncovers early developmental enhancers in humans. Nature 470, 279–83. doi:10.1038/nature09692

Ramanan, N., Shen, Y., Sarsfield, S., Lemberger, T., Schütz, G., Linden, D.J., Ginty, D.D., 2005. SRF mediates activity-induced gene expression and synaptic plasticity but not neuronal viability. Nat. Neurosci. 8, 759–767. doi:10.1038/nn1462

Ramirez-Carrozzi, V.R., Braas, D., Bhatt, D.M., Cheng, C.S., Hong, C., Doty, K.R., Black, J.C., Hoffmann, A., Carey, M., Smale, S.T., 2009. A Unifying Model for the Selective Regulation of Inducible Transcription by CpG Islands and Nucleosome Remodeling. Cell 138, 114–128. doi:10.1016/j.cell.2009.04.020

Ramirez-Carrozzi, V.R., Nazarian, A.A., Li, C.C., Gore, S.L., Sridharan, R., Imbalzano, A.N., Smale, S.T., 2006. Selective and antagonistic functions of SWI/SNF and Mi-2B nucleosome remodeling complexes during an inflammatory response. Genes Dev. 20, 282–296. doi:10.1101/gad.1383206

Saha, R.N., Dudek, S.M., 2013. Splitting Hares and Tortoises: A classification of neuronal immediate early gene transcription based on poised RNA polymerase II. Neuroscience. doi:10.1016/j.neuroscience.2013.04.064

Saha, R.N., Wissink, E.M., Bailey, E.R., Zhao, M., Fargo, D.C., Hwang, J.-Y., Daigle, K.R., Fenn, J.D., Adelman, K., Dudek, S.M., 2011. Rapid activity-induced transcription of Arc and other IEGs relies on poised RNA polymerase II. Nat. Neurosci. 14, 848–56. doi:10.1038/nn.2839

Santos, S.D.M., Verveer, P.J., Bastiaens, P.I.H., 2007. Growth factor-induced MAPK network topology shapes Erk response determining PC-12 cell fate. Nat. Cell Biol. 9, 324–30. doi:10.1038/ncb1543

Schwanhausser, B., Busse, D., Li, N., Dittmar, G., Schuchhardt, J., Wolf, J., Chen, W., Selbach, M., 2011. Global quantification of mammalian gene expression control. Nature 473, 337–342. doi:10.1038/nature10098

Sgambato, V., Vanhoutte, P., Pagès, C., Rogard, M., Hipskind, R., Besson, M.J., Caboche, J., 1998. In vivo expression and regulation of Elk-1, a target of the extracellular-regulated kinase signaling pathway, in the adult rat brain. J. Neurosci. 18, 214–226.

Sharrocks, A.D., 1995. ERK2/p42 MAP kinase stimulates both autonomous and SRF-dependent DNA binding by Elk-1. FEBS Lett. 368, 77–80. doi:10.1016/0014-5793(95)00604-8

Sheng, H.Z., Fields, R.D., Nelson, P.G., 1993. Specific Regulation of Immediate Early Genes by Patterned Neuronal Activity. J. Neurosci. Res. 35, 459–467.

Shi, C., Davis, M., 2001. Visual and pain pathways involved in fear conditioning measured with fear-potentiated startle: Behavioral and anatomic studies. J. Neurosci. 21, 9844–9855. doi:10.1109/ICONIP.2002.1202138

Spiegel, I., Mardinly, A.R., Gabel, H.W., Bazinet, J.E., Couch, C.H., Tzeng, C.P., Harmin, D.A., Greenberg, M.E., 2014. Npas4 regulates excitatory-inhibitory balance within neural circuits through cell-type-specific gene programs. Cell 157, 1216–29. doi:10.1016/j.cell.2014.03.058

Su, Y., Shin, J., Zhong, C., Wang, S., Roychowdhury, P., Lim, J., Kim, D., Ming, G., Song, H., 2017. Neuronal activity modifies the chromatin accessibility landscape in the adult brain. Nat. Neurosci. 20. doi:10.1038/nn.4494

Sullivan, J.M., Badimon, A., Schaefer, U., Ayata, P., Gray, J., Chung, C.-W., von Schimmelmann, M., Zhang, F., Garton, N., Smithers, N., Lewis, H., Tarakhovsky, A., Prinjha, R.K., Schaefer, A., 2015. Autism-like syndrome is induced by pharmacological suppression of BET proteins in young mice. J. Exp. Med. 212, 1771–81. doi:10.1084/jem.20151271

Szutorisz, H., Dillon, N., Tora, L., 2005. The role of enhancers as centres for general transcription factor recruitment. Trends Biochem. Sci. 30, 593–599. doi:10.1016/j.tibs.2005.08.006

Telese, F., Ma, Q., Perez, P.M., Notani, D., Oh, S., Li, W., Comoletti, D., Ohgi, K.A., Taylor, H., Rosenfeld, M.G., 2015. LRP8-Reelin-Regulated Neuronal Enhancer Signature Underlying Learning and Memory Formation. Neuron 1–15. doi:10.1016/j.neuron.2015.03.033

Thomas, G.M., Huganir, R.L., 2004. MAPK cascade signalling and synaptic plasticity. Nat. Rev. Neurosci. 5, 173–83. doi:10.1038/nrn1346

Toettcher, J.E., Weiner, O.D., Lim, W. a, 2013. Using optogenetics to interrogate the dynamic control of signal transmission by the Ras/Erk module. Cell 155, 1422–34. doi:10.1016/j.cell.2013.11.004

Treisman, R., 1996. Regulation of transcription by MAP kinase cascades. Curr. Opin. Cell Biol. 8, 205–215. doi:10.1016/S0955-0674(96)80067-6

Tullai, J.W., Schaffer, M.E., Mullenbrock, S., Sholder, G., Kasif, S., Cooper, G.M., 2007. Immediate-early and delayed primary response genes are distinct in function and genomic architecture. J. Biol. Chem. 282, 23981–23995. doi:10.1074/jbc.M702044200

Turrigiano, G.G., 2008. The self-tuning neuron: synaptic scaling of excitatory synapses. Cell 135, 422–35. doi:10.1016/j.cell.2008.10.008

Wang, G., Balamotis, M.A., Stevens, J.L., Yamaguchi, Y., Handa, H., Berk, A.J., 2005. Mediator requirement for both recruitment and postrecruitment steps in transcription initiation. Mol. Cell 17, 683–694. doi:10.1016/j.molcel.2005.02.010

West, A.E., Greenberg, M.E., 2011. Neuronal Activity–Regulated Gene Transcription in Synapse Development and Cognitive Function 1–21.

Worley, P.F., Bhat, R. V, Baraban, J.M., Erickson, A., Mcnaughton, L., 1993. Thresholds for Synaptic Activation of Transcription Factors Hippocampus: Correlation with Long-term Enhancement in. J. Neurosci. 73.

Wu, G., Deisseroth, K., Tsien, R.W., 2001. Spaced stimuli stabilize MAPK pathway activation and its effects on dendritic morphology. Nat. Neurosci. 151–158.

Wu, G., Deisseroth, K., Tsien, R.W., 2000. Activity-dependent CREB phosphorylation: Convergence of a fast, sensitive calmodulin kinase pathway and a slow, less sensitive mitogen-activated protein kinase pathway. Proc. Natl. Acad. Sci.

Xia, Z., Dudek, H., Miranti, C.K., Greenberg, M.E., 1996. Calcium Influx via the NMDA Receptor Induces Immediate Early Gene Transcription by a MAP Kinase / ERK-Dependent Mechanism. J. Neurosci. 16, 5425–5436.

Yamamoto, K.R., Alberts, B.M., 1976. Steroid Receptors: Elements for modulation of eukaryotic transcription. Annu. Rev. Biochem. 721–746. doi:10.1007/978-1-4939-1346-6

Zhai, S., Ark, E.D., Parra-Bueno, P., Yasuda, R., 2013. Long-distance integration of nuclear ERK signaling triggered by activation of a few dendritic spines. Science 342, 1107—11. doi:10.1126/science.1245622

Zheng, F., Luo, Y., Wang, H., 2009. Regulation of BDNF-mediated transcription of immediate early gene Arc by intracellular calcium and calmodulin. J Neurosci Res 87, 380—392. doi:10.1002/jnr.21863.Regulation

Zhu, Y., Sun, L., Chen, Z., Whitaker, J.W., Wang, T., Wang, W., 2013. Predicting enhancer transcription and activity from chromatin modifications. Nucleic Acids Res. 41, 10032–10043. doi:10.1093/nar/gkt826

## Methods References

Dale, R.K., Matzat, L.H., Lei, E.P., 2014. Metaseq: A Python package for integrative genome-wide analysis reveals relationships between chromatin insulators and associated nuclear mRNA. Nucleic Acids Res. 42, 9158–9170. doi:10.1093/nar/gku644

Dobin, A., Davis, C.A., Schlesinger, F., Drenkow, J., Zaleski, C., Jha, S., Batut, P., Chaisson, M., Gingeras, T.R., 2013. STAR: Ultrafast universal RNA-seq aligner. Bioinformatics 29, 15–21. doi:10.1093/bioinformatics/bts635

Hunter, J.D., 2007. Matplotlib: A 2D graphics environment. Comput. Sci. Eng. 9, 99–104. doi:10.1109/MCSE.2007.55

Kuhn, R.M., Haussler, D., James, Kent W., 2013. The UCSC genome browser and associated tools. Brief. Bioinform. 14, 144–161. doi:10.1093/bib/bbs038

Langmead, B., Salzberg, S.L., 2012. Fast gapped-read alignment with Bowtie 2. Nat Methods 9, 357–359. doi:10.1038/nmeth.1923

Li, H., Handsaker, B., Wysoker, A., Fennell, T., Ruan, J., Homer, N., Marth, G., Abecasis, G., Durbin, R., 2009. The Sequence Alignment/Map format and SAMtools. Bioinformatics 25, 2078–2079. doi:10.1093/bioinformatics/btp352

Quinlan, A.R., Hall, I.M., 2010. BEDTools: A flexible suite of utilities for comparing genomic features. Bioinformatics 26, 841–842. doi:10.1093/bioinformatics/btq033

Robinson, M.D., McCarthy, D.J., Smyth, G.K., 2009. edgeR: A Bioconductor package for differential expression analysis of digital gene expression data. Bioinformatics 26, 139–140. doi:10.1093/bioinformatics/btp616

Saha, R.N., Ghosh, A., Palencia, C.A., Fung, Y.K., Dudek, S.M., Pahan, K., 2009. TNF-alpha preconditioning protects neurons via neuron-specific up-regulation of CREB-binding protein. J. Immunol. (Baltimore, Md 1950) 183, 2068–2078. doi:10.4049/jimmunol.0801892

Shao, Z., Zhang, Y., Yuan, G.-C., Orkin, S.H., Waxman, D.J., 2012. MAnorm: a robust model for quantitative comparison of ChIP-Seq data sets. Genome Biol. 13, R16. doi:10.1186/gb-2012-13-3-r16

Van Der Walt, S., Colbert, S.C., Varoquaux, G., 2011. The NumPy array: A structure for efficient numerical computation. Comput. Sci. Eng. 13, 22–30. doi:10.1109/MCSE.2011.37

